# Comparative analysis of CRISPR/Cas9-targeted nanopore sequencing approaches in repeat expansion disorders

**DOI:** 10.1101/2024.12.04.626786

**Authors:** Louise Benarroch, Pierre-Yves Boëlle, Hélène Madry, Badreddine Mohand Oumoussa, Nobuyuki Eura, Ichizo Nishino, Karim Labrèche, Valeriia Gorbunova, Guillaume Bassez, Tanya Stojkovic, Geneviève Gourdon, Gisèle Bonne, Stéphanie Tomé

## Abstract

More than 50 repeat expansion disorders have been identified, with long-read sequencing marking a new milestone in the diagnosis of these disorders. Despite these major achievements, the comprehensive characterization of short tandem repeats in a pathological context remains challenging, primarily due to their inherent characteristics such as motif complexity, high GC content, and variable length.

In this study, our aim was to thoroughly characterize repeat expansions in two neuromuscular diseases: myotonic dystrophy type 1 (DM1) and oculopharyngodistal myopathy (OPDM) using CRISPR/Cas9- targeted long-read sequencing (Oxford Nanopore Technologies, ONT). We conducted precise analyses of the DM1 and OPDM loci, determining repeat size, repeat length distribution, expansion architecture and DNA methylation, using three different basecalling strategies (MinKnow software, Dorado and Bonito). We demonstrated the importance of the basecalling strategy in repeat expansion characterization. We proposed guidelines to perform CRISPR-Cas9 targeted long-read sequencing (no longer supported by ONT), from library preparation to bioinformatical analyses. Finally, we showed, for the first time, somatic mosaicism, hypermethylation of *LRP12* loci in OPDM symptomatic patients and changes in the repeat tract structure of these patients.

We propose a strategy based on CRISPR/Cas9-enrichment long-read sequencing for repeat expansion diseases, which could be readily applicable in research but also in diagnostic settings.

## INTRODUCTION

Over the last 30 years, more than 50 repeat expansion disorders (RED) have been identified. Fragile X syndrome (FXS, OMIM #300624) and spinal and bulbar muscular atrophy (SBMA, OMIM #313200) were the first RED identified caused by a CGG expansion in the 5′-untranslated region (5’-UTR) of *FMR1* gene [1–4] and a CAG expansion in the coding region of *AR* gene [5], respectively. Since then, the exponential development of genetic tools and next generation sequencing technologies such as long-read sequencing (LRS), mark a new milestone in RED diagnosis with the discovery of new disorders over the last decade [6]. However, fully characterizing these expansions (number of motifs, expansion architecture, methylation status, somatic mosaicism, etc.) remains challenging due to their inherent complexity. In this study, we focused on two disorders, myotonic dystrophy type 1 (DM1) and oculopharyngodistal myopathy (OPDM). DM1 is the most common adult muscular dystrophy, characterized by multisystemic symptoms among which muscle weakness, myotonia, cataracts, cardiac conduction defects, respiratory distress and neurological impairments. DM1 is caused by a highly unstable CTG expansion in the 3’-UTR of *DMPK* gene and is one of the most complex REDs due to genetic anticipation, age-dependent and tissue-specific somatic instability [7]. OPDM is a rare neuromuscular disorder characterized by progressive ptosis, ophthalmoplegia, dysphagia and facial and distal limb weakness. OPDM is due to a CGG repeat expansion in the 5’-UTR of five different genes identified so far: *LRP12* (OPDM1) [8]*, GIPC1* (OPDM2) [9], *NOTCH2NLC* (OPDM3) [10], *RILPL1* (OPDM4) [11], and *ABCD3* gene (OPDM5) [12].

Despite the great effort to decipher the genetic cause of rare genetic disorders, only one third of them have been solved using short-read sequencing. The low diagnostic rate can be explained in part by the mutation localization within complex regions, inaccessible by short-read sequencing approaches such as short tandem repeat regions [13]. LRS, now, give access to those regions, opening a new era for the diagnosis of rare genetic diseases, especially for REDs. Two LRS approaches are available provided by Pacific Bioscience (PacBio) and Oxford Nanopore Technologies (ONT) which are slowly introduced into the diagnostic field. Here, we used ONT’s long-read approaches, particularly the targeted sequencing approach using CRISPR/Cas9 strategy, to assess repeat expansions at DM1 and OPDM loci. Despite available protocols provided by ONT and open access bio-informatic pipelines, we realized that precisely characterizing expansion in our cases remained challenging. Indeed, inconsistency in sequencing metrics (i.e. read length distribution, coverage, etc.) as well as sequencing errors were prone to appear depending on the library preparation protocol used and/or the basecalling strategies apply to our raw data. The choice of the basecaller (BC), to translate the changes of the raw electrical signal into a nucleotidic sequence appeared to be critical when investigating tandem repeats. Here, we tested different applications available on ONT: CRISPR/Cas9-enrichment, adaptive sampling (AS) and whole genome-LRS (WG) in 5 DM1 patients and 4 OPDM patients, in total. We then tested the performance for different basecalling strategies, the MinKnow software implemented in PromethION sequencer and two standalone open-sourced BC, Dorado and Bonito. And finally, we investigated the architecture of the repeat expansion by assessing their structure and their methylation levels.

## RESULTS

### Generation of sequencing files

Libraries were sequenced using PromethION device implemented with the MinKnow software allowing to perform real-time analysis of the electrical signal. Until October 2023, MinKnow used the Guppy BC after which it changed to Dorado BC. Fast5 files (containing raw electrical signal) were converted into FastQ files either using MinKnow, or with the open-sourced BC Dorado (v5.3) and Bonito (v7.2). Frequent updates of MinKnow throughout this study (See Materials and methods section, Table 5) prevented us from using it to characterize repeat expansion in our sample set. We therefore used the standalone version of Dorado BC (v5.3) to determine sequencing metrics, compare performance to MinKnow software and Bonito BC performances and finally investigate repeat expansions in DM1 and OPDM patient samples.

### CRISPR/Cas9 long-read sequencing metrics of DM1 and OPDM loci

We performed LRS using ONT technologies on DNA extracted from 9 patients, 5 DM1 patients and 4 OPDM patients (Table 1). We compared the 2 available chemistries at the time, ONT R9.4 and ONT R10.4 to perform CRISPR/Cas9-based LRS to target specifically the DM1 locus (*DMPK* gene) and 3 OPDM loci (*LRP12*, *GIPC1* and *NOTCH2NLC* genes). Basecalling was performed using Dorado BC (v5.3). Coverage was estimated using the number of total reads and their phred score. Phred score is widely used to estimate the accuracy of the calling for each base, and we can interpret this score as an estimation of error. For instance, a phred score of 20 (Q20) estimates 1% chance of error, so 99% of accuracy. In sequencing data, Q20 or above is considered sufficient and we have used that score as a quality cut-off throughout this study. Using R9.4 chemistry, we observed a mean phred score of 17.8 [range: 15.3-21.9], across all reads. On average, less than 40% of reads reached Q20 cut-off, regardless of the disease investigated. Chemistry R10.4 significantly improved reads quality, as expected, with a mean quality score of 29.9 [27.3-31] and more than 70% of total reads reached Q20 score (Mann-Whitney test, average quality [9.4] *vs* [10.4] p-value = 0.0012) (Table 1). The R10.4 chemistry also provided a similar phred score (around 30) for both WG and AS. Interestingly, the expanded allele average quality for all reads (AvgQ all) was lower than the normal allele with R10.4 chemistry, suggesting an impact of the repeat expansion on sequencing quality (Mann-Whitney test, average quality of [normal allele] *vs* [expanded allele] p-value = 0.0411) (Table 2). Regarding WG and AS, no significant changes in average quality was observed between both alleles. However, with only one patient tested and a low number of reads, we cannot make a definite conclusion on the impact of the expansion in these applications.

**Table 1.**
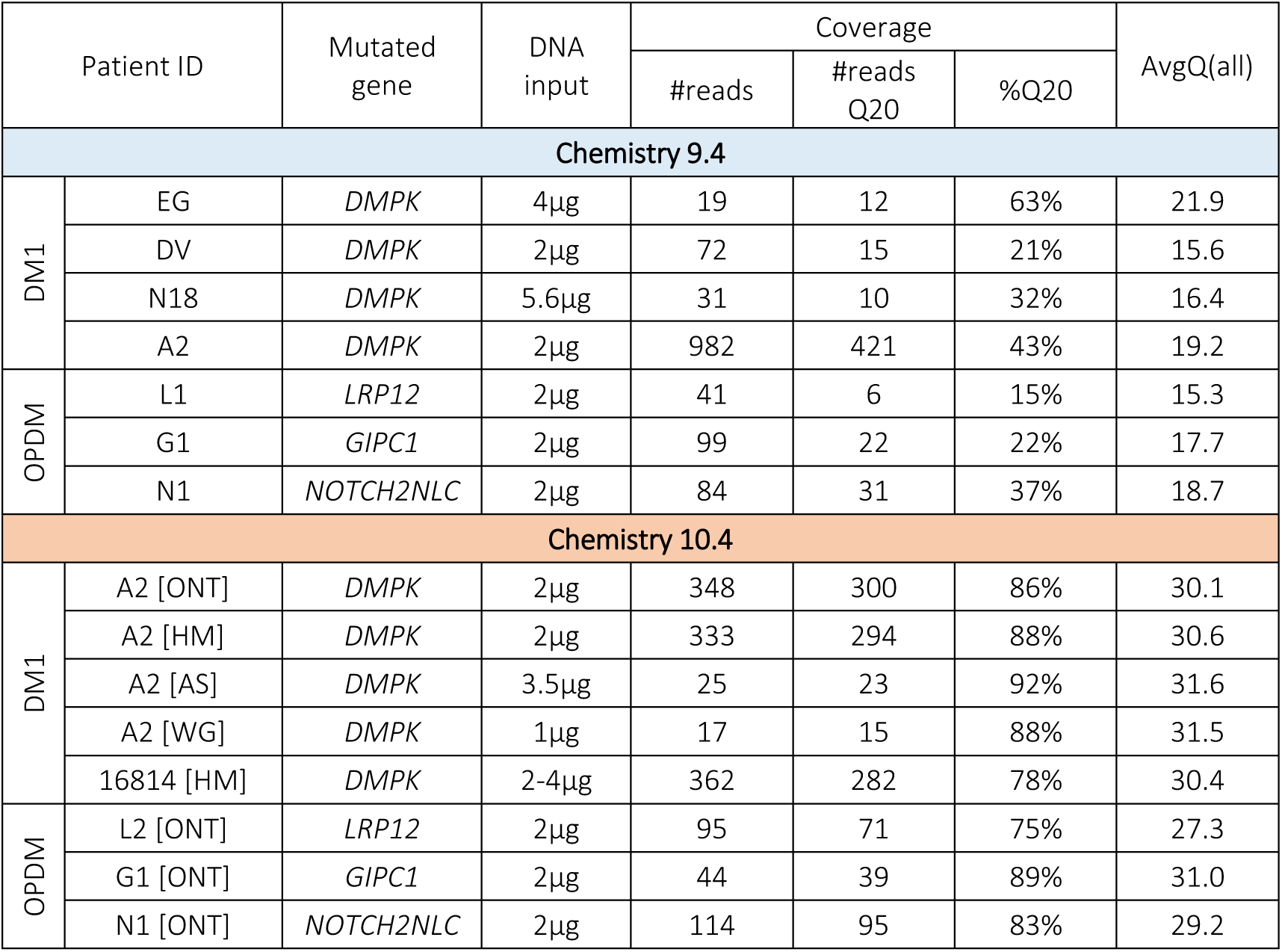
Long-read sequencing metrics for DM1 and OPDM patients. All reads were analyzed using Dorado BC (v5.3). Coverage was determined by the number of total reads (#reads), number of reads with a mapping quality above 20 (#reads Q20) and the percentage of reads with a mapping quality above 20 (%Q20). Average quality was assessed for all reads [AvgQ(all)] using a phred-scale quality score. In chemistry R10.4, different protocols were used: ONT protocol [ONT], home-made protocol [HM], adaptive sampling [AS] and whole genome sequencing [WG].

| Patient ID |  | Mutated gene | DNA input | Coverage |  |  | AvgQ(all) |
| --- | --- | --- | --- | --- | --- | --- | --- |
|  |  |  |  | #reads | #reads Q20 | %Q20 |  |
| Chemistry 9.4 |  |  |  |  |  |  |  |
| DM1 | EG | DMPK | 4µg | 19 | 12 | 63% | 21.9 |
|  | DV | DMPK | 2µg | 72 | 15 | 21% | 15.6 |
|  | N18 | DMPK | 5.6µg | 31 | 10 | 32% | 16.4 |
|  | A2 | DMPK | 2µg | 982 | 421 | 43% | 19.2 |
| OPDM | L1 | LRP12 | 2µg | 41 | 6 | 15% | 15.3 |
|  | G1 | GIPC1 | 2µg | 99 | 22 | 22% | 17.7 |
|  | N1 | NOTCH2NLC | 2µg | 84 | 31 | 37% | 18.7 |
| Chemistry 10.4 |  |  |  |  |  |  |  |
| DM1 | A2 [ONT] | DMPK | 2µg | 348 | 300 | 86% | 30.1 |
|  | A2 [HM] | DMPK | 2µg | 333 | 294 | 88% | 30.6 |
|  | A2 [AS] | DMPK | 3.5µg | 25 | 23 | 92% | 31.6 |
|  | A2 [WG] | DMPK | 1µg | 17 | 15 | 88% | 31.5 |
|  | 16814 [HM] | DMPK | 2-4µg | 362 | 282 | 78% | 30.4 |
| OPDM | L2 [ONT] | LRP12 | 2µg | 95 | 71 | 75% | 27.3 |
|  | G1 [ONT] | GIPC1 | 2µg | 44 | 39 | 89% | 31.0 |
|  | N1 [ONT] | NOTCH2NLC | 2µg | 114 | 95 | 83% | 29.2 |

**Table 2.**
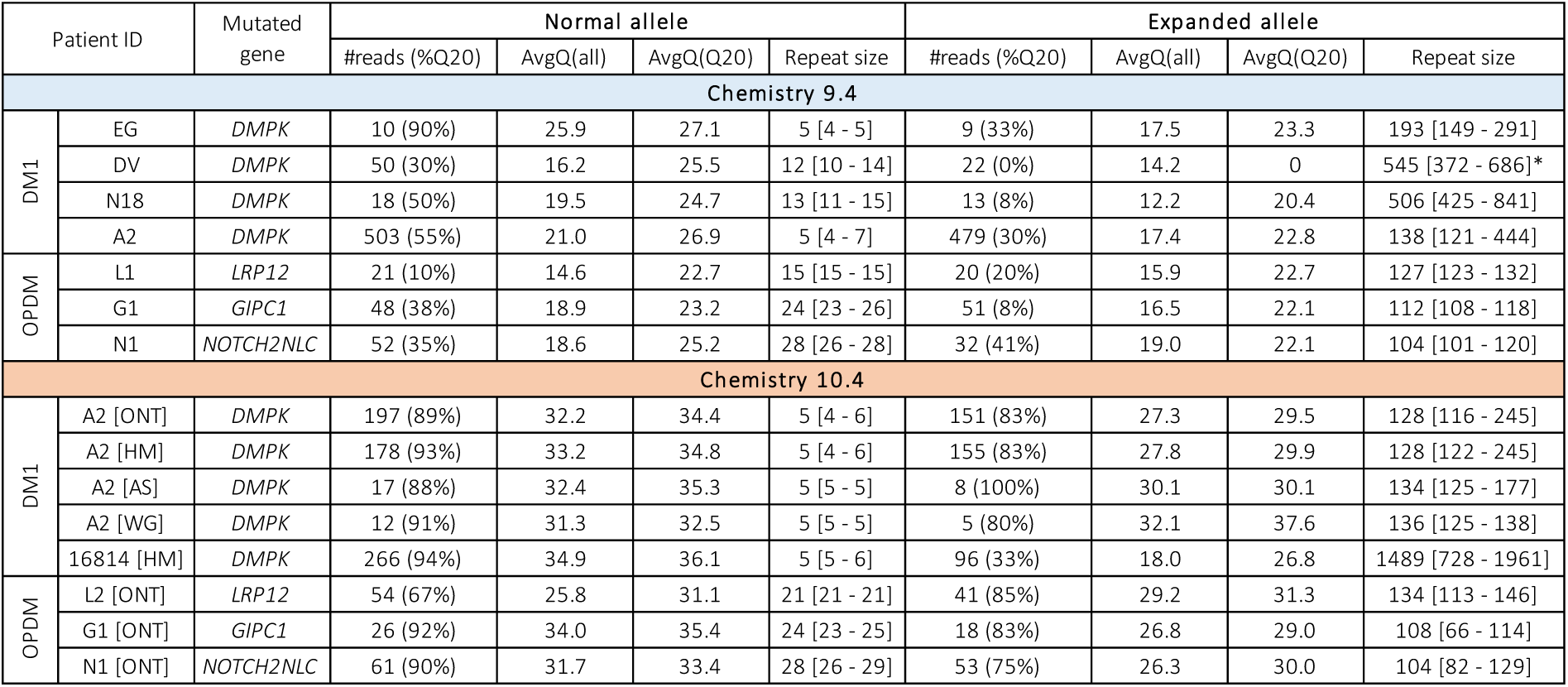
Long-read sequencing metrics on the normal and expanded alleles in DM1 and OPDM patients. All reads were analyzed using Dorado BC (v5.3). For both allele, coverage is expressed as the total number of reads (#reads) with the percentage of Q20 reads (%Q20). Average quality (Avg Q) was assessed by calculating the mean quality for all reads [AvgQ(all)] or for reads with a phred score above 20 [AvgQ(Q20)] specific of each allele. Repeat size for the normal and expanded allele are defined by the median [min – max] of Q20 reads. Repeat size marked by “*” were estimated using all reads as no reads reached the quality cut-off of 20. The estimated median size for the A2 patient varied slightly between approaches due to differences in the number of molecules analyzed, but all methods consistently indicated ∼130 repeats. ONT protocol [ONT], home-made protocol [HM], adaptive sampling [AS] and whole genome sequencing [WG].

| Patient ID |  | Mutated gene | Normal allele |  |  |  | Expanded allele |  |  |  |
| --- | --- | --- | --- | --- | --- | --- | --- | --- | --- | --- |
|  |  |  | #reads (%Q20) | AvgQ(all) | AvgQ(Q20) | Repeat size | #reads (%Q20) | AvgQ(all) | AvgQ(Q20) | Repeat size |
| Chemistry 9.4 |  |  |  |  |  |  |  |  |  |  |
| DM1 | EG | DMPK | 10 (90%) | 25.9 | 27.1 | 5 [4 - 5] | 9 (33%) | 17.5 | 23.3 | 193 [149 - 291] |
|  | DV | DMPK | 50 (30%) | 16.2 | 25.5 | 12 [10 - 14] | 22 (0%) | 14.2 | 0 | 545 [372 - 686]* |
|  | N18 | DMPK | 18 (50%) | 19.5 | 24.7 | 13 [11 - 15] | 13 (8%) | 12.2 | 20.4 | 506 [425 - 841] |
|  | A2 | DMPK | 503 (55%) | 21.0 | 26.9 | 5 [4 - 7] | 479 (30%) | 17.4 | 22.8 | 138 [121 - 444] |
| OPDM | L1 | LRP12 | 21 (10%) | 14.6 | 22.7 | 15 [15 - 15] | 20 (20%) | 15.9 | 22.7 | 127 [123 - 132] |
|  | G1 | GIPC1 | 48 (38%) | 18.9 | 23.2 | 24 [23 - 26] | 51 (8%) | 16.5 | 22.1 | 112 [108 - 118] |
|  | N1 | NOTCH2NLC | 52 (35%) | 18.6 | 25.2 | 28 [26 - 28] | 32 (41%) | 19.0 | 22.1 | 104 [101 - 120] |
| Chemistry 10.4 |  |  |  |  |  |  |  |  |  |  |
| DM1 | A2 [ONT] | DMPK | 197 (89%) | 32.2 | 34.4 | 5 [4 - 6] | 151 (83%) | 27.3 | 29.5 | 128 [116 - 245] |
|  | A2 [HM] | DMPK | 178 (93%) | 33.2 | 34.8 | 5 [4 - 6] | 155 (83%) | 27.8 | 29.9 | 128 [122 - 245] |
|  | A2 [AS] | DMPK | 17 (88%) | 32.4 | 35.3 | 5 [5 - 5] | 8 (100%) | 30.1 | 30.1 | 134 [125 - 177] |
|  | A2 [WG] | DMPK | 12 (91%) | 31.3 | 32.5 | 5 [5 - 5] | 5 (80%) | 32.1 | 37.6 | 136 [125 - 138] |
|  | 16814 [HM] | DMPK | 266 (94%) | 34.9 | 36.1 | 5 [5 - 6] | 96 (33%) | 18.0 | 26.8 | 1489 [728 - 1961] |
| OPDM | L2 [ONT] | LRP12 | 54 (67%) | 25.8 | 31.1 | 21 [21 - 21] | 41 (85%) | 29.2 | 31.3 | 134 [113 - 146] |
|  | G1 [ONT] | GIPC1 | 26 (92%) | 34.0 | 35.4 | 24 [23 - 25] | 18 (83%) | 26.8 | 29.0 | 108 [66 - 114] |
|  | N1 [ONT] | NOTCH2NLC | 61 (90%) | 31.7 | 33.4 | 28 [26 - 29] | 53 (75%) | 26.3 | 30.0 | 104 [82 - 129] |

Unfortunately, CRISPR/Cas9-enrichment protocol is no longer supported by ONT. Therefore, we decided to set up a ‘home-made’ protocol using similar existing products combined with the adapter ligation steps of ONT ligation sequencing kit (see Materials and Methods section). We tested our protocol on patient A2, which protocol achieved similar results to the ONT protocol in terms of coverage, percentage of Q20 reads and average quality (Table 1 and 2). Our in-house protocol was also validated in patient #16814, who carries more than 1000 CTG repeats.

### Basecaller comparison on average quality of the expanded allele’s reads

Several BC are available to convert raw electrical signals into nucleotidic sequences. MinKnow, software developed by ONT, is commonly used for real-time analysis and basecalling. Over the course of our study (as mentioned before), MinKnow initially used the Guppy BC and then transitioned to Dorado. In addition to MinKnow, a standalone version of Dorado and another open-sourced BC, Bonito were also available. Both Dorado “standalone” and Bonito provided free and user-friendly alternatives to MinKnow. Therefore, we decided to compare the basecalling accuracy of MinKnow with that of Dorado (v5.3) and Bonito (v7.2). We used data from patients analyzed with R10.4 chemistry as we saw a significant increase in reads’ quality compared to R9.4 chemistry, independently of the BC and the pathology (Fig. 1a).

**Figure 1.**
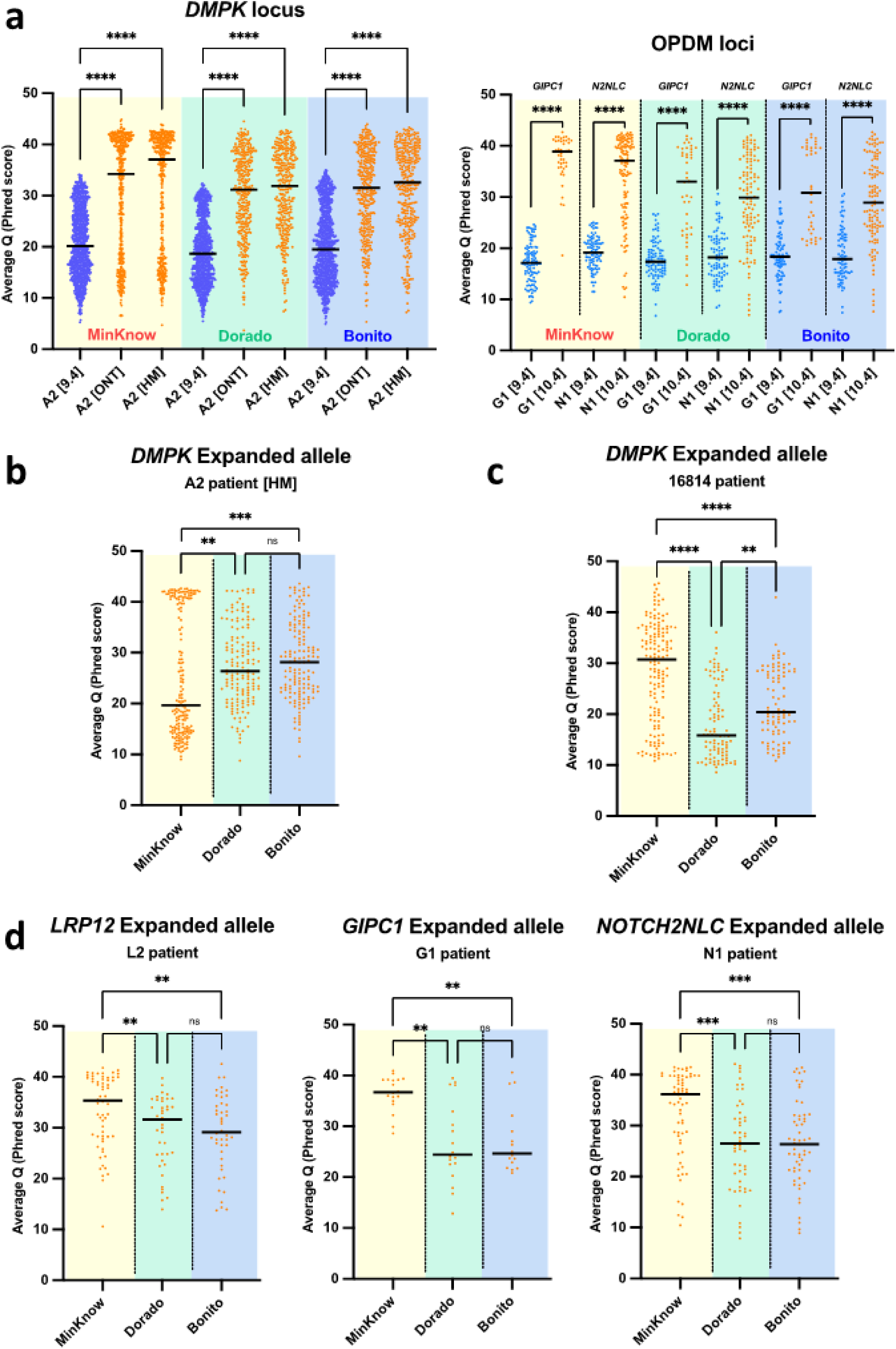
Basecallers comparison on reads’ average quality at DM1 and OPDM loci. **(a)** Comparison of average quality of all reads between R9.4 and R10.4 chemistry at DM1 (left panel) and OPDM (right panel) loci. For all graphs, ONT R9.4 chemistry is represented with blue dots, ONT R10.4 chemistry (home-made ‘HM’) with orange dots. Average quality was estimated with MinKnow (yellow frame), Dorado (green frame) and Bonito (blue frame). [N2NLC – NOTCH2NLC]. **(b-d)** Expanded allele average quality using R10.4 chemistry for **(b)** A2 patient, **(c)** #16814 patient and **(d)** OPDM patients, L2 (left panel), G1 (middle panel) and N1 (right panel). Kruskal-Wallis test, ns – not significant; [*] p-value < 0.05; [**] p-value < 0.01; [***] p-value < 0.001 and [****] p-value < 0.0001

By focusing on A2 patient and depending on the approach used, WG or targeted approaches (CRISPR/Cas9-enrichment or AS), BC performances differed regarding the expanded allele (Table 3). In the targeted approaches (CRISPR/Cas9 and AS), we show that Dorado and Bonito exhibited overall better performances than MinKnow. More specifically, for the CRISPR/Cas9 strategy, A2 [HM] average quality for MinKnow was lower than Dorado and Bonito, reaching 25.9 *vs* 27.8 and 28.7, respectively (Kruskal-Wallis test, MinKnow *vs* Dorado p-value = 0.0075, MinKnow *vs* Bonito p-value = 0.0001) (Fig. 1b, Table 3). In addition, for adaptive sampling, A2 [AS] average quality for MinKnow was also lower than Dorado and Bonito, with 13.7 *vs* 30.1 and 30.9 (Kruskal-Wallis test, MinKnow *vs* Dorado p-value = 0.0126, MinKnow *vs* Bonito p-value 0.0063) (Table 3). However, when applying a Q20 quality cut-off for the expanded allele in A2 [HM], MinKnow outperformed Dorado and Bonito, with an average quality score of 37.6 compared to 29.9 and 30.6, respectively. Despite this higher average score with MinKnow, the number of Q20 reads remained higher when using Dorado or Bonito BC, trend also observed in AS (Table 3). Regarding WG, MinKnow had an average quality score higher than the other two BC (A2 [WG] average quality MinKnow *vs* Dorado and Bonito, 34.8 *vs* 32.1 and 28.4) but the differences were not significant (Kruskal-Wallis test, A2 [WG] MinKnow *vs* Dorado p-value > 0.99, MinKnow *vs* Bonito p-value > 0.99) (Table 3). It is important to note that, for the normal allele, MinKnow consistently produced higher quality scores than Dorado and Bonito, except in the case of AS (Fig. S1a, Table S1). To complement our data on BC comparison in DM1, we also analyzed a second DM1 patient, #16814, who carries a longer expansion than A2 patient (1,489 *vs* 128 repeats) (Table 2). For patient #16814, MinKnow produced a higher average quality score for both the normal and expanded alleles, which could be explained by either the update of MinKnow software or by the fact that MinKnow may be better suited for analyzing longer expansions compared to Dorado and Bonito (Fig. 1c, Fig. S1a, Table S1).

**Table 3.** Basecaller comparison on average quality of the expanded allele. Three basecallers were tested: MinKnow (yellow), Dorado (green) and Bonito (blue). Average quality was assessed by calculating the mean quality for all reads [AvgQ(all)] or for reads with a phred score above 20 [AvgQ(Q20)]. AvgQ - Average quality (phred score); %Q20 - percentage of Q20 reads on total expanded allele reads.

| Exp. allele<br>(10.4) | Sample ID | MinKnow |  |  | Dorado |  |  | Bonito |  |  |
| --- | --- | --- | --- | --- | --- | --- | --- | --- | --- | --- |
|  |  | AvgQ (all) | %Q20 | AvgQ (Q20) | AvgQ (all) | %Q20 | AvgQ (Q20) | AvgQ (all) | %Q20 | AvgQ (Q20) |
| DMPK | A2 [ONT] | 24.9 | 49% | 35.5 | 27.3 | 83% | 29.5 | 28.5 | 89% | 29.9 |
|  | A2 [HM] | 25.9 | 48% | 37.6 | 27.8 | 83% | 29.9 | 28.7 | 87% | 30.6 |
|  | A2 [AS] | 13.7 | 0% | 0 | 30.1 | 100% | 30.1 | 30.9 | 100% | 30.9 |
|  | A2 [WG] | 34.8 | 91% | 37.2 | 32.1 | 80% | 37.6 | 28.4 | 63% | 37.4 |
|  | 16814 [HM] | 28.4 | 74% | 33.2 | 18.0 | 33% | 26.8 | 22.1 | 54% | 27.1 |
| LRP12 | L2 [ONT] | 33.3 | 97% | 34.0 | 29.2 | 85% | 31.3 | 29.0 | 86% | 31.1 |
| GIPC1 | G1 [ONT] | 36.5 | 100% | 36.5 | 26.8 | 83% | 29.0 | 27.3 | 100% | 27.3 |
| NOTCH2NLC | N1 [ONT] | 32.5 | 91% | 34.4 | 26.3 | 75% | 30.0 | 26.5 | 79% | 29.5 |

For all three OPDM patients (L2, G1 and N1), MinKnow displayed a better average quality score regardless of the allele analyzed (phred score > 30) (Fig. 1d, Fig. S1b, Table 3 and Table S1). For L2 patient, reads targeting *LRP12* expanded allele displayed a higher average score with MinKnow than Dorado and Bonito (Kruskal-Wallis test, MinKnow *vs* Dorado, p-value = 0.0083; MinKnow *vs* Bonito, p-value = 0.0063, Dorado *vs* Bonito, p-value > 0.99) (Fig. 1d, Table 3). A similar observation is made for G1 patient, where the expansion is in the *GIPC1* gene (Kruskal-Wallis test, MinKnow *vs* Dorado, p-value = 0.0011; MinKnow *vs* Bonito, p-value = 0.0021; Dorado *vs* Bonito, p-value > 0.99) (Fig. 1d, Table 3). For N1 patient, reads targeting the expanded allele of *NOTCH2NLC* gene also displayed a significantly higher average quality score with MinKnow than with Dorado and Bonito (Kruskal-Wallis test, [10.4] MinKnow *vs* Dorado, p-value = 0.0004; MinKnow *vs* Bonito, p-value = 0.0004; Dorado *vs* Bonito, p-value > 0.99) (Fig. 1d). Similar results were obtained for the normal allele for *LRP12* and *NOTCH2NLC* genes (Fig. S1b). For G1 patient, no significant difference was observed between MinKnow and Bonito (Fig. S1b). Similar results were observed for each patient when focusing on Q20 reads, with MinKnow providing better average quality compared to the other BC (Table 3).

By comparing DM1 A2 [ONT] and OPDM L2 [ONT] patients, and thus the type of codon that are expanded (CTG *vs* CGG), L2 patient’s reads showed better quality scores than those targeting the *DMPK* gene when using MinKnow (Mann-Whitney test, MinKnow average quality A2 [ONT] *vs* L2 [ONT], p-value < 0.0001). This difference is lost when comparing Dorado (Mann-Whitney test, Dorado average quality A2 [ONT] *vs* L2 [ONT], p-value = 0.13) or Bonito quality scores (Mann-Whitney test, Bonito average quality A2 [ONT] *vs* L2 [ONT], p-value = 0.52). Our data suggested that the type of repeat investigated can also have an impact on BC performances.

Based on our results and despite a small set of patients, we could show that the choice of BC is a critical step on LRS data analysis and should be adapted regarding sequencing approach used (targeted *vs* WG), size of the expansion and the type of triplet repeat. We recommend testing at least two BC, MinKnow and Dorado, for each sequencing experiments. Despite a better performance of MinKnow, the inconsistencies in the BC versions implemented in the software (see Materials and Methods section, Table 5) prevented us to use MinKnow data to compare our samples. Therefore, from now on, our analyses are obtained using Dorado “standalone” (v5.3) BC on Q20 reads.

### Estimation of repeat expansions length and somatic mosaicism

DM1 A2 patient and other family members have been extensively investigated and well characterized using different sequencing techniques: classical PCR, 3′ triplet-primed-PCR (TP-PCR), Single-Molecule Real Time (SMRT) long read sequencing (Pacific Bioscience) and now ONT-LRS [14–16]. In summary, all techniques confirmed the presence of the expansion in these patients, estimated its size (around 150 repeats), identified a single CAG interruption within the expanded region and revealed a low level of somatic mosaicism compared to size- and age-matched classical DM1 patients. Therefore, from here on out, A2 [HM] patient will be used as an experimental control to determine repeat expansion length, somatic mosaicism, and the presence of interruptions. The repeat size frequency distribution showed that this patient carried 5 CTG repeats (± 1 tract due to sequencing error) on the normal allele and between 122 and 245 motifs in the expanded allele, with a median at 128, highlighting the somatic mosaicism already described in this patient (Fig. 2a, Table 2). For #16814 DM1 patient, the normal allele carried 5 CTG repeats (± 1 tract due to sequencing error) and between 728 and 1,961 CTG repeats, with a median at 1,489 (Fig. 2b, Table 2). The great variability in CTG repeat range on the expanded allele highlighted somatic mosaicism with repeat expansion in #16814 being more unstable than repeat expansion in A2 patient (Fig. S2). For OPDM patients, the repeat size frequency distribution of the normal allele median was at 21, 24 and 28 for *LRP12*, *GIPC1* and *NOTCH2NLC* genes (± 1 tract due to sequencing error), respectively (Fig. 2c-e). Regarding the expanded allele, the median was at 134, 108 and 104 for *LRP12*, *GIPC1* and *NOTCH2NLC* genes, respectively. Repeat sizes ranged from 113 to 146 for *LRP12*, from 66 to 114 for *GIPC1* and 82 to 129 for *NOTCH2NLC*, highlighting for the first time, somatic mosaicism in OPDM, independently of the analyzed locus (Fig. 2c-e, Table 2). However, further investigations in a larger cohort of patients are required to fully attest the somatic mosaicism, which is not the scope of this paper.

**Figure 2.**
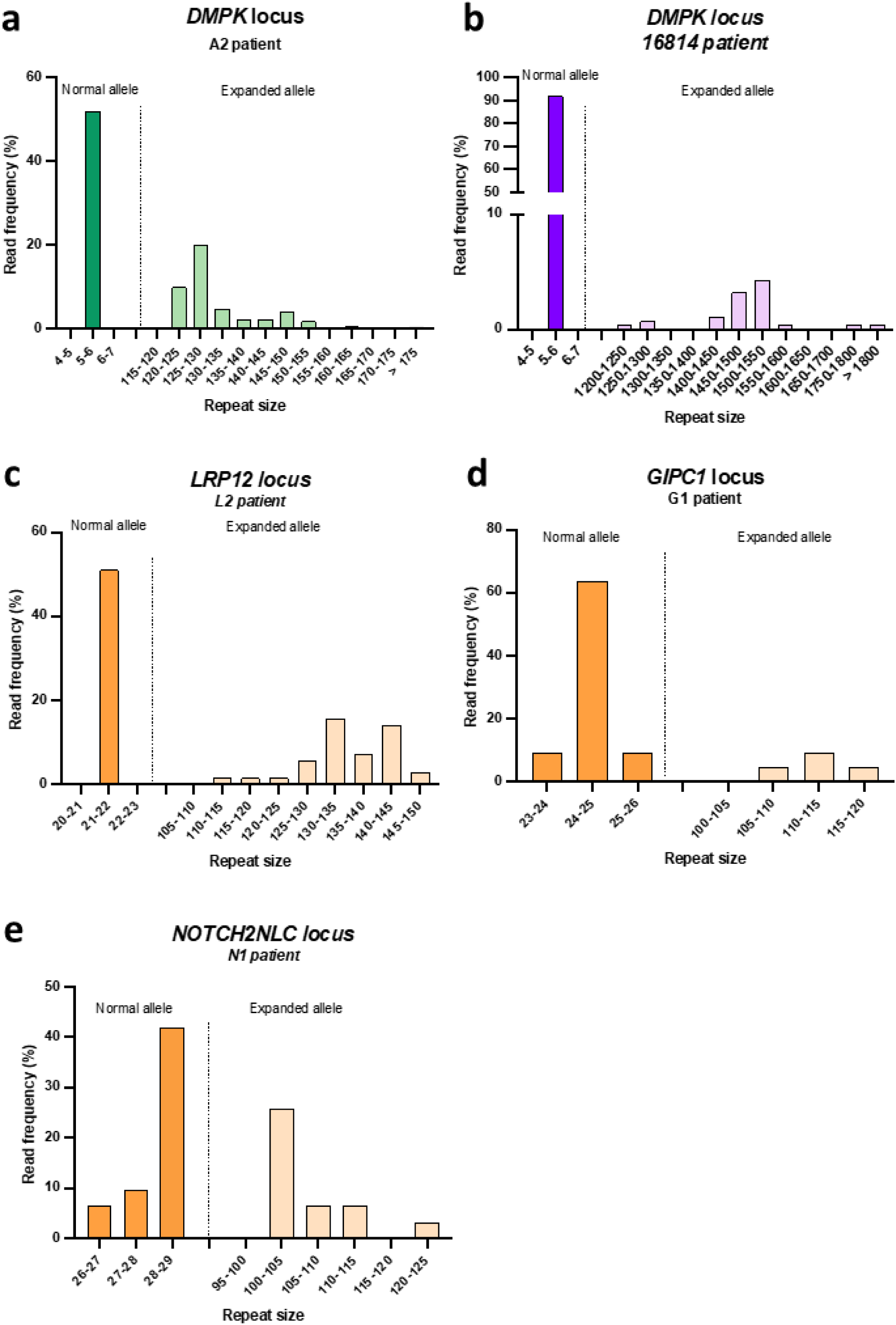
Read length distribution at **(a-b)** DM1 and **(c-e)** OPDM loci. Read length was estimated using Q20 reads. The x-axis displays the length of the repeats detected (number of repeats, 3bp per repeat) for each locus: **(a-b)** DMPK, **(c)** LRP12, **(d)** GIPC1, and **(e)** NOTCH2NLC. The normal and expanded alleles are separated by a striated line. The y-axis displays read frequency calculated as a ratio of number of Q20 reads on total number of reads.

### Changes in expansion sequence’s architecture in OPDM patients

Using CRISPR/Cas9-targeted LRS (R10.4 chemistry), a single CAG interruption in the expanded allele of A2 [HM] patient was clearly identified as previously described [14] (Fig. 3a). This result validates ONT-LRS suitable for clearly determining sequence architecture changes in trinucleotide repeat diseases. In all samples, we observed reads containing artefactual motifs due to sequencing artefacts (Fig. S3) which were largely eliminated by selecting only reads with a phred score above 20 (Fig. 3). Patient #16814 displayed a pure CTG expansion without any artefactual interruptions (Fig. 3b). The expanded region containing around 1,500 repeats, it is more prone for error to occur and a phred score above 20 was not sufficient to eliminate of all sequencing inaccuracies. But choosing a quality cut-off above 30 for this patient excluded most of the reads targeting the expanded allele. At *LRP12* loci (patient L2), reads were aligned using a TAG motif located right before the start of the CCG repeat tract in the reference sequence (File S1). In the normal allele, the repeat sequence is framed by two motifs, 5’- (ACG)2-CCG-ACG-3’ at the 5’ end and 5’-AGC-CAC-CGG-3’ at the 3’ end of the repeats. In the expanded allele, those two motifs are lost, replaced by only one ACG codon at the 5’ end of the expansion and a single ‘G’ at the 3’ end of the expansion (Fig. 3c and File S1). The loss of those motifs was also found in another *LRP12* mutated patient (L1 patient, File S1), supporting the fact that changes in *LRP12* sequence structure is a consequence of the presence of the expansion. At *GIPC1* locus (G1 patient), reads were aligned using CCC motif, upstream of the GCC expansion. In the normal allele, the repeat sequence started with 5’-TCC-GCC-TCC-3’ motif in the reference sequence that is lost in the expanded allele and replaced by 5’-(TCC)4-GCC-(GCG)3-3’ interruption (Fig. 3d and File S1). Finally, *NOTCH2NLC* (N1 patient) reads were aligned using CCA motif, upstream of the GGC expansion. In the normal allele, GGC repeats ended with 5’-(GGA)2-(GGC)2-3’ motif whereas in the expanded allele, additional GGC are inserted between the two GGA, creating a 5’-GGA-(GGC)3-GGA-(GGC)5-3’ motif (Fig. 3e and File S1).

**Figure 3.**
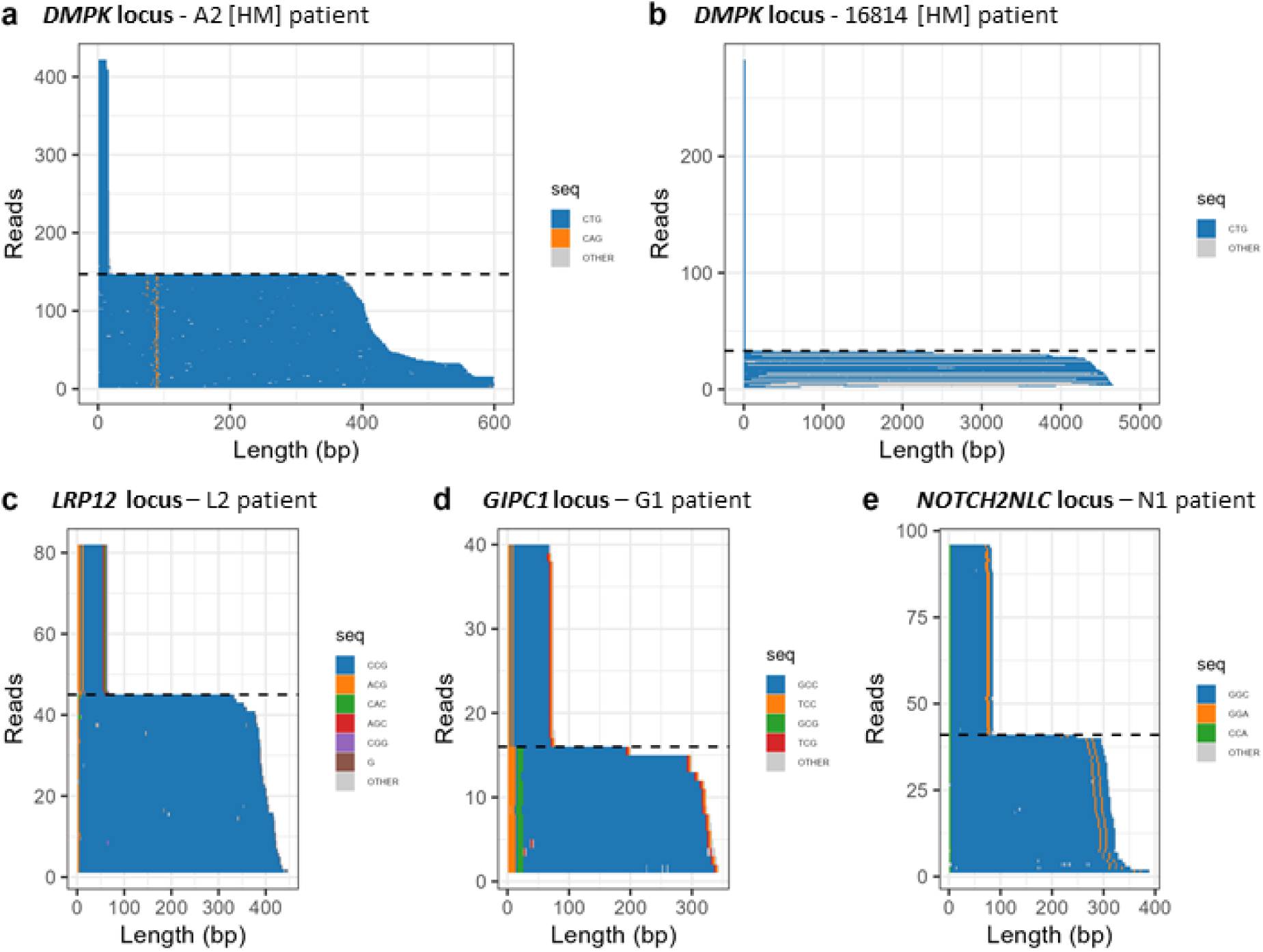
Waterfall plots highlighting the expansion structures at **(a-b)** DM1 and **(c-e)** OPDM loci. Waterfalls were generated using Dorado BC (v5.3) on Q20 reads. For each waterfall, the x-axis shows the number of repeats (in base pairs) detected in each read. The y-axis displays each individual read ranked by repeat length. The plateau reflects multiple reads carrying the same length, distinguishing the normal allele and the expanded allele. The normal allele is above the striated line whereas the expanded allele is under the striated lines. The reference codon **(a-b)** CTG, **(c)** CCG, **(d)** GCC or **(e)** GGC is represented in blue; different colours are used to highlight codons within the repeat sequence that are not the reference codon, within the expansion.

Thus, CRISPR/Cas9-targeted sequencing allowed us to highlight somatic mosaicism in OPDM but also to identify changes in the structure of repeated tracts in the three OPDM genes, never described before : *i)* in *LRP12* expansion, loss of two complex motifs associated with the expansion; *ii)* in *GIPC1* expansion, presence of a 5’-(TCC)4-GCC-(GCG)3-3’ interruption; and *iii)* in *NOTCH2NLC* expansion, presence of 5’-GGA-(GGC)3-GGA-(GGC)5-3’ interruptions.

### DNA methylation profiling at DM1 and OPDM loci

To assess DNA methylation pattern, we used Dorado BC (v5.3) and Modkit, a methylation-calling pipeline to detect DNA modifications not only within the flanking region but also within the expanded region. Several DNA modifications can be identified using ONT-LRS such as 5-methylcytosine (5mC), 5-hydroxymethylcytosine (5hmC) and N6-methyladenine (6mA), where 5mC is the most common methylation mark. To perform 5mC analyses, we first used WG data from the A2 patient to analyze the *CNBP* gene methylation profile, as it is known to be hypermethylated in the genomic region immediately downstream of the CCTG repeats in DM2 [17] and DM1 patients (unpublished data). We confirmed the hypermethylation of *CNBP* gene in A2 [HM] patient, supporting that methylation profiling of the 5mC mark can be performed using ONT-LRS (Fig. 4a). We did not see any changes in 5mC methylation profile either within the expanded region or in the flanking regions for A2 patient (Fig. 4b), and OPDM patients, G1 and N1 (Fig. S4a-b). However, we observed hypermethylation in the *LRP12* locus (Fig. 4c). Indeed, in L2 patient, we detected an increase in methylation frequency in *LRP12* 5’UTR region, reaching approximatively 10% hypermethylation that was present in the flanking region and within the expanded region (Fig. 4d). An increase in methylation frequency was also observed in another *LRP12* patient (L1 patient) that was sequenced using the R9.4 chemistry confirming the hypermethylation in the flanking regions as well as within the expanded region (Fig. S4c-d). We can conclude that CGG expansion caused changes in methylation profile of *LRP12* locus. To our knowledge it is the first report of an existing hypermethylation in a symptomatic OPDM patient.

**Figure 4.**
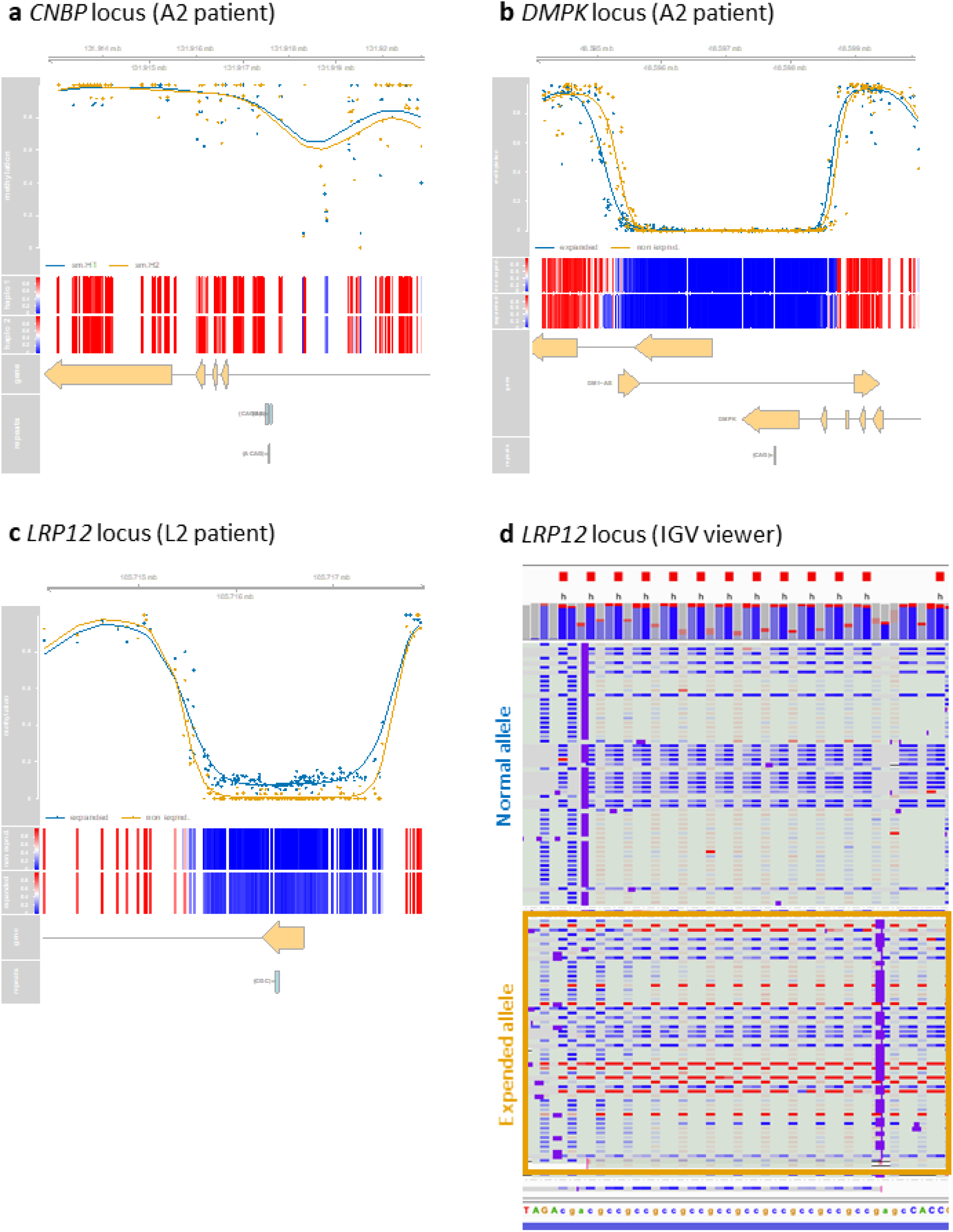
5-methylcytosine (5mC) methylation levels at **(a)** CNBP, **(b)** DMPK and **(c)** LRP12 loci. Each graph displays, from top to bottom, the genomic localization; the methylation levels where each dot represent one read on the normal allele (yellow) or the expanded allele (blue); heatmap of methylation levels; gene(s) located in the region of interest; and localization of the repeats. **(a)** CNBP allele separation was performed using SNPs (Single Nucleotide Polymorphism) and sequence-level differences between the two alleles, as no expansion could allow us to distinguish alleles based on its size. **(b-c)** Allele separation was performed using the size of triplet repeated region. **(d)** IGV screenshot of the LRP12 expanded region, displaying the methylation status of each read. The normal allele (top) and the expanded allele (bottom) are separated by a dotted line. Each line represents one read. Unmethylated sites are colored in blue, methylated sites are colored in red. The expanded region is surrounded by a yellow frame.

The 5hmC modification is a product of the demethylation of 5mC-methylated site but also a novel epigenetic regulator, however its functions have yet to be clearly established. It has been described to play a role in several diseases but has never been studied in DM1 and OPDM. In this study we observed that 5hmC profiles are similar between normal and expanded alleles at the DM1 and OPDM loci (Fig. S5). However, we cannot rule out that the absence of signal is due to the incapacity of the ONT to detect such mark in our data. ONT also enabled the detection of 6mA modifications at the DM1 and OPDM loci. In all patient tested, we observed a high variability in 6mA frequency compared to the 5mC frequency which is more homogeneous. (Fig. 5a-e). The methylation frequency appeared to be quite extensive around the 3’-UTR region of *DMPK* (Fig. 5a-b) and high around the 5’-UTR region of the three OPDM genes, *LRP12* (Fig. 5c), *NOTCH2NLC* (Fig. 5d) and *GIPC1* (Fig. 5e). Although some variations are observed between normal and expanded alleles for *LRP12* and *GIPC1*, it is difficult to conclude due to the high variability of the 6mA profiles, especially without a positive control for this mark.

**Figure 5.**
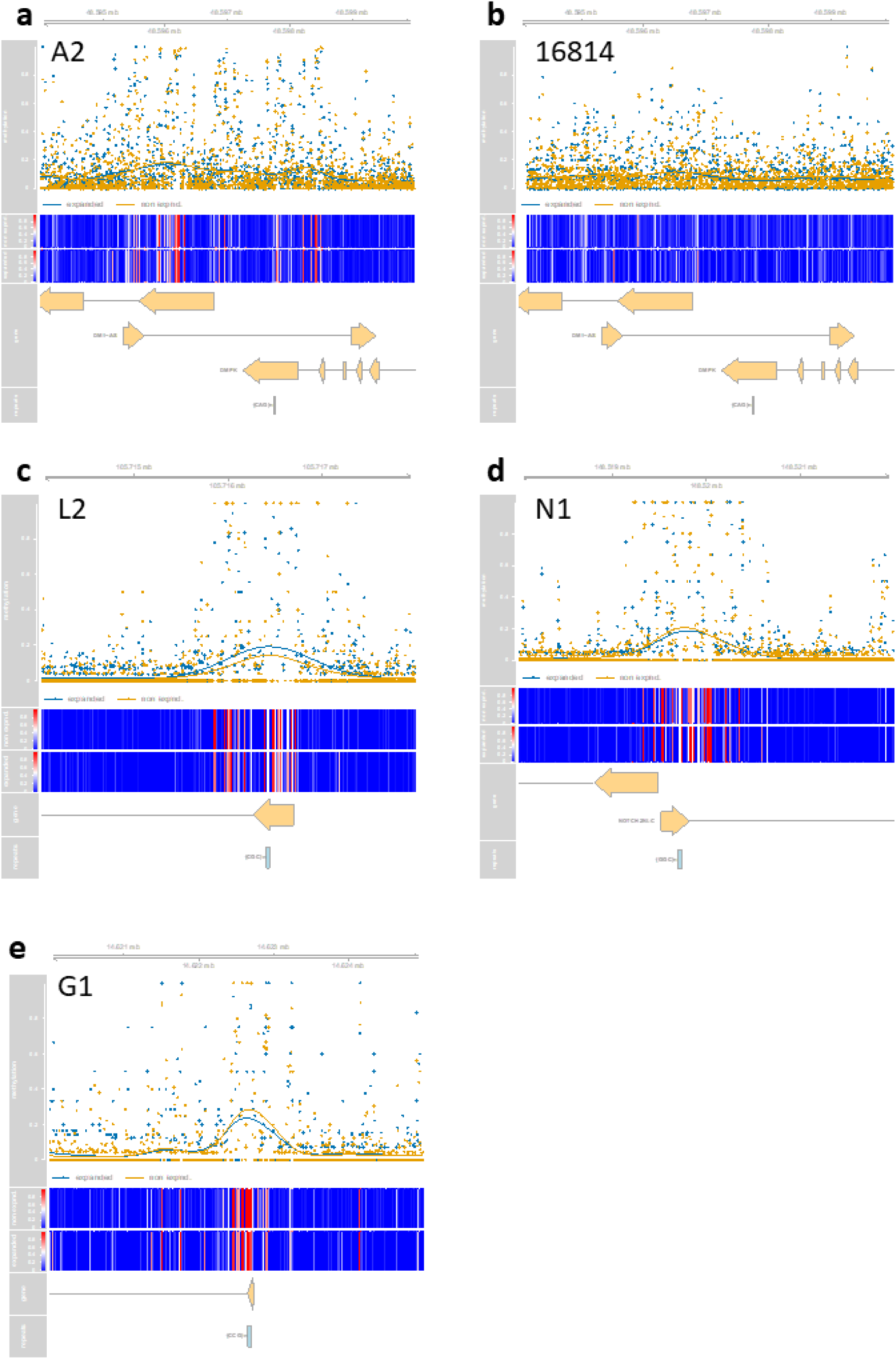
N6-methyldeoxyadenosine (6mA) methylation levels at **(a-b)** DM1 and **(c-e)** OPDM loci. Each graph displays, from top to bottom, the genomic localization; the methylation levels where each dot represent one read on the normal allele (yellow) or the expanded allele (blue); heatmap of methylation levels; gene(s) located in the region of interest; and localization of the repeats.

## DISCUSSION

In this study we showed that LRS is a suitable strategy to investigate REDs but requires some adjustments depending on the disease of interest. Thanks to ONT’s multiple sequencing protocols (including WG, AS, and CRISPR/Cas9-targeted sequencing), we showed that ONT-LRS can *i)* detect the presence of repeat expansions, *ii)* estimate the size and structure of the repeat tract and *iii)* determine the 5mC methylation pattern. This was possible because, in our rather small cohort (9 patients in total), we included one DM1 patient (A2) as a control, who had been thoroughly characterized using multiple approaches (classical PCR, TP-PCR, PacBio-LRS). By investigating A2 patients, we confirmed what was shown with gold-standard approaches: median expansion of 130 CTG triplets interrupted by a CAG triplet at the 30^th^ position, along with a low level of somatic mosaicism. Therefore, additional patients and validation using conventional approaches, as performed for A2 patient, will be necessary to *i)* confirm the presence of somatic mosaicism in OPDM patients and *ii)* accurately define the methylation profiles for 5hmC and 6mA marks. Finally, we propose guidelines for DM1 and OPDM expanded loci characterization. Our results showed that the R10.4 chemistry significantly improved sequencing metrics in expanded regions for both DM1 and OPDM patients. We compared two protocols for CRISPR/Cas9 library preparation (ONT provided and home-made) and we did not see any significant differences between them. So, we propose here a suitable alternative to continue CRISPR/Cas9- enrichment using the R10.4 chemistry, in a research setting. ONT developed another targeted approach called adaptive sampling. Similarly to CRISPR/Cas9-enrichement, AS enable the sequencing of a region of interest (ROI) but it can select and sequence only reads mapping to the ROI in real-time by rejecting off-target reads. AS was presented as more efficient, less time-consuming and cost-effective compared to CRISPR/Cas9-enrichment. Surprisingly, AS did not provide better coverage or average quality that we could expect, compared to CRISPR/Cas9 protocol using DNA from the same extraction. However, low coverage (less than 20X) prevented us to make any definite conclusion regarding the pros and cons of AS compared to CRISPR/Cas9 approach. Nevertheless, CRISPR/Cas9 enrichment offers key advantages over AS: *i)* it allows multiplexing (2–5 samples per flow cell *vs*. one for AS), and *ii)* it generates smaller datasets (<10 Gb *vs*. AS >20 Gb), reducing sequencing time, cost, and data storage needs. These factors should be considered when choosing between CRISPR/Cas9 and AS. While ONT- LRS shows considerable promise in addressing the diagnostic challenges associated with genetic diseases [18], its application on REDs in a diagnostic setting still requires improvements. Particularly, there are notable variations observed between flow cells, impacting pore availability, coverage or multiplexing (data not shown) and the bioinformatics tools. These inconsistencies represent significant obstacles to ensure the reliability and reproducibility of LRS-based diagnostics for REDs. Optimization of our LRS protocol will be necessary for future applications to decrease the technical variability we observed. Different aspect of the protocol may be improved such as the quality of the genomic DNA, fragmentation of the DNA prior to library preparation as well as an additional purification step to eliminate the unrequired DNA fragments.

Basecalling is also a critical step in the sequencing workflow as it translates the electrical signal received by the pore into the DNA sequence, hence all downstream investigations depend on it. ONT used 2 BC implemented in its MinKnow software: Guppy followed by Dorado since October 2023. ONT provided frequent updates of the BC implemented in MinKnow software. They performed several updates of Dorado BC since its implementation in MinKnow. Although frequent updates improve basecalling performance, they can pose challenges when comparing data obtained at different time points during long-term studies, as it was the case in our study, or when using such a pipeline in a diagnostic setting. It is important to note that frequent updates are also available for Dorado “standalone” as 12 new versions has been released from February 2024 (Dorado v5.3) to January 2025 (Dorado v9.1). However, unlike MinKnow software, we can retain older versions of the BC and compare data between them. Therefore, to overcome inconsistency in data analysis, we decided to use the standalone version of Dorado (v5.3) to compare our samples, enabling us to draw any conclusion on repeat expansion detection in our settings. Hence, we could conclude that the choice of BC should depend on the type of study that will be conducted (short- *vs* long-term), the sequencing approach (targeted *vs* WG), the size of the expansion and the type of the repeated motif (CTG *vs* CGG). Our study highlighted the importance of appropriate BC and emphasized the need to consider both the software version, and the specific neural network used in bioinformatics analyses, as these evolve rapidly over time. Indeed, we observed discrepancies in sequencing data that may be due to differences in each BC’s neural network to translate the electrical signal of CTG *vs* CGG triplet where the process remains unclear in the analysis. However, by analyzing the indels between MinKnow and Dorado (v5.3), we observed that small indels and the substitutions in reads with lesser quality (phred score < 30) disappeared in high quality read (phred score

> 30). One possible explanation is that the electrical signal from one particular triplet is close to the signal from another. Therefore, to precisely determine the number of repeats, the architecture of the expanded allele and flanking regions as well as the presence of somatic mosaicism, our data suggest that BC selection is crucial and the combination of several of them may also be needed in some cases [19]. We strongly recommend evaluating several BC performances before making any conclusions [20,21]. Additionally, the details of the filters used and the importance of the phred quality score are crucial for each ONT analysis, as highlighted in our study. We focused on the Dorado BC (v5.3) allowing us to effectively compare our samples to analyze somatic mosaicism, the nucleotidic sequence structure of expansions, and DNA methylation.

ONT allowed us to determine not only the size but also the structure of the repeat tracts which can both have a huge impact on patients’ prognosis and the clinical variability of the disease. For instance, the presence of trinucleotide interruption within the repeat tract can cause different pathophysiological mechanisms underlying disease development and progression. In different REDs, interruptions have been thoroughly investigated, and these studies showed the biological impact of these interruptions, either protective or aggravating, in disease clinical manifestations or progression. In fragile X syndrome (FXS), AGG codons that commonly interrupt the CGG repeat expansion in normal individuals, are less frequent in FXS patients causing transmission instability, hence increasing the expansion risk. The presence of AGG interruption, within the expansion, act as a stabilizing factor preventing instability of the repeats. Knowing the AGG interruption size can help estimate the risk for FXS intermediate individuals (45 to 54 CGG repeats) or pre-mutated individual (55 to 200 repeats) to develop a full-mutation expansion (over 200 repeats) [22]. In DM1 patients, the presence of CCG, CTC or CGG interruptions within the CTG expansion can either be associated with additional atypical neurological symptoms or can prevent or delay severe onset of the disease [23]. These interruptions as well as an unique CAG interruption have been associated with the stabilization of the repeat through unknown mechanisms [14,23–30]. In OPDM patients, changes in the expansion sequence have never been described before, to our knowledge. However, it has been shown that in other *NOTCH2NLC*-associated diseases, GGA interruption within the GGC repeat tract can cause different phenotypes based on the number of repeats, going from Parkinson disease (less than 100 repeats) to dominant muscle weakness (over 200 repeats) [31]. In our *NOTCH2NLC*-mutated patient (N1 patient), we did not find any GGA expansion as in the other *NOTCH2NLC*-associated disease but a change in the sequence motif with the insertion of GGC codons between the two GGA. However, more *NOTCH2NLC*-associated OPDM patients would be required to confirm this motif, and to attest if it is specific to OPDM patients carrying *NOTCH2NLC* mutation and investigate its potential role in the pathophysiological mechanisms of the disease. Regarding *LRP12* and *GIPC1* expansions, no loss of motif or presence of interruption have ever been reported, but as for *NOTCH2NLC* patients, additional patients would be required to confirm these observations.

One of ONT’s strength is the direct detection of DNA or RNA modifications, during the translocation of the molecule through the pore. Already available with R9.4 chemistry, R10.4 chemistry allowed us to detect three types of modification: 5-methylcytosine (5mC), its oxidative counterpart 5- hydroxymethylcytosine (5hmC) and N6-methyldeoxyadenine (6mA), in the flanking regions as well as within the expanded region. The 5mC mark is the most abundant, representing 70-80% of cytosine preceding a guanidine (CpG) mainly located in promoter regions, into CpG islands [32]. DNA methylation changes were observed in many diseases, among which OPDM. Interestingly in OPDM, the epigenetic changes were only described in asymptomatic patients carrying longer expansion than the disease-causing range, in *GIPC1*, *NOTCH2NLC, RILPL1 and ABCD3* genes [9,11,33–35]. As expected, we did not see any changes in affected OPDM patients carrying either a *GIPC1* or *NOTCH2NLC* expansion. However, we notice an increase in 5mC methylation levels (around 10%) in *LRP12* patients that has never been reported before. The use of a positive control for 5mC detection with *CNBP* gene hypermethylation allowed us to firmly conclude that the hypermethylation is not a sequencing artefact. Moreover, we observed a hypermethylation of *LRP12* expanded region in another patients, supporting this observation. We hypothesized that due to the expansion of a CCG codon in OPDM patients, it may increase the methylation levels of the expanded region in affected individuals but not sufficiently enough to have a protective impact, as the one observed in asymptomatic carriers. Similar increase was observed in affected *NOTCH2NLC* patients (around 30% of methylation) but the difference with healthy individuals was not significant and was associated with an increase mRNA level in muscle but not in blood, supporting our hypothesis [36]. To go further we investigated the 5hmC modification, a product of the demethylation of 5mC-methylated. It is considered as a novel epigenetic regulator, but its functions have yet to be clearly established [37]. We did not see any changes in 5hmC levels in any patients which was not surprising as it is lowly found in mammalian tissues. Indeed, 5hmC content reach approximatively 0.1%, with the highest being in the brain with 1% [37]. However, as we did not have a positive control for 5hmC detection, we could not rule out a technical limitation of ONT on 5hmC detection. Finally, to our knowledge it is the first time that 6mA methylation have been investigated in REDs. Preferentially found in prokaryote, 6mA modification represent 0.05% of the total adenines in the human genome and are found preferentially in the brain, as 5hmC modifications. This mark is enriched in coding region of transcriptionally active genes and mostly occur on [G/C]AGG[C/T] motif [38]. In our study, we observed great variability of the methylation frequency at the different loci, but the frequency did not seem to be significantly impacted by the presence of the expansion. Contrary to what is described in the literature [38], the frequency of 6mA modifications seems higher in the 5’-UTR region that in the intronic region, especially for OPDM loci. However, so little is known about 6mA modification that is still too early to make any definite conclusions on if what we observed is relevant and regarding the role of these modifications in OPDM and DM1.

## CONCLUSION

Despite the small cohort of DM1 and OPDM patients analyzed in this study, the findings highlight the importance of sequencing and bioinformatical strategies when studying REDs. We provide guidelines and recommendations to investigate DM1 and OPDM loci that could hopefully help in the diagnostic and the care of patients.

## METHODS

### DNA samples

High molecular weight DNA from DM1 patients’ blood or lymphoblastoid cell lines were extracted either using Nanobind CBB HMW DNA extraction from cells, bacteria and blood kit (Pacific Bioscience, catalog n°102-207-600, Menlo Park, CA) or using Chemagic 360 system (Revvity chemagen Technologie Gmbh, Bernolsheim, France). DNA samples from OPDM patients were kindly provided by Pr. Ichizo Nishino’s group at the National Center of Neurology and Psychiatry (Tokyo, Japan). DNA were extracted from blood samples of four patients carrying expansion either in *LRP12* gene (L1 and L2 patients), *GIPC1* gene (G1 patient) or *NOTCH2NLC* gene (N1 patient). For all DNA samples, quality and quantity were assessed using Qubit fluorometer (Thermofisher Scientific, catalog n°Q33238, Asnières-sur-Seine, France) and the TapeStation 4150 System (Agilent Technologies, catalog N°G2992AA, Les Ulis, France). All DNA samples showed a predominant fragment size greater than 50 kb, which is considered suitable for long-read sequencing.

### Oxford Nanopore long-read sequencing

#### CRISPR/Cas9-enrichment

Targeted long-read sequencing was performed on DNA samples from both DM1 and OPDM patients, in a singleplex and/or multiplex setting. Three different protocols were tested, two from ONT: “Ligation Sequencing gDNA-Cas9 enrichment protocol” (Chemistry R9.4; SQK- CS9109 version: CAS_9106_v109_revD_16Sep2020) and “Ligation sequencing kit V14” (Chemistry R10.4; SQK-LSK114 version: GDE_9161_v114_revT_29Jun2022), and one “home-made” protocol (Chemistry R10.4). For all three protocols, dephosphorylation of genomic DNA (gDNA), Cas9 cleavage and dA-tailing were performed following the Cas9 enrichment protocol (ONT, catalog n°SQK-CS9109 – discontinued in March 2024, Paris, France). In brief, 2-5μg of gDNA was dephosphorylated, to prevent Cas9 off-target. In parallel, Cas9-ribonucleoprotein complexes (Cas9-RNPs) were formed by pooling 100μM of each Alt-R CRISPR/Cas9 crRNA (Integrated DNA Technologies – IDT, Leuven, Belgium) specific of each targeted loci (*LRP12*-F: 5’-TCTTTACCTCTTACGGCAAG-3’; *LRP12*-R: 5’- TCTGGTCCCGTAGAGTCCAC-3’; *GIPC1*-F: 5’-GACGCTGTTGTCATTACTAT-3’; *GIPC1*-R: 5’- ACATCTAAGCTGCTCACCAT-3’; *NOTCH2NLC*-F: 5’-GGAGACATCTTGGTGCATAC-3’; *NOTCH2NLC*-R: 5’-AAGCGCATTTGTACATGAGA-3’; *DMPK*-5mc-US1: 5’- ACCCAAGGCTCGCCCAATA-3’; *DMPK*-5mc-US2: 5’-GGGGAGAGCGGTACCACTTG-3’; *DMPK*-5mc-DS1: 5’-GACGAGGTTACTTCAGACAT-3’; *DMPK*-5mc-DS2 [39]: 5’-GACCTGCGAGTCACACAAC-3’), 50 μM transactivation crRNA (tracrRNA, IDT) and 62μM Alt-R S.p. HiFi Cas9 Nuclease V3 (IDT, catalog n°1081061). CRISPR/Cas9 crRNA targeting *HTT* gene were also used as a positive control (data not shown), according to the manufacturer’s recommendations: *HTT*-F1: 5’-TTTGCCCATTGGTTAGAAGC-3’; *HTT*-R1: 5’-GGACAAAGTTAGGTACTCAG-3’; *HTT*-F2: 5’- TCTTATGAGTCTGCCCACTG-3’; *HTT*-R2: 5’-CTAGACTCTTAACTCGCTTG-3’.

The dephosphorylated gDNA was then incubated with Cas9-RNPs, allowing Cas9 to cleave at the different loci. Non-templated A nucleotides (dATP) were added to gDNA blunt ends formed by Cas9 cleavage using TAQ polymerase, adding a dA-tail to prepare DNA for adapter ligation and sequencing. The adapter ligation steps differ slightly when using the Cas9 sequencing kit V109 or the ligation sequencing kit V114, the main difference being the adapter mix used, Adapter Mix (AMX) or Ligation Adapter (LA), respectively. Adapters were added to the DNA thanks to the T4 ligase, either provided by the ligation sequencing kit or by the blunt/TA ligase master mix (New England Biolabs – NEB, catalog n°M0367S, Evry, France). End-prepped DNA were purified using AMPure XP Beads (Beckman Coulter, catalog n°A63880, Villepinte, France) and eluted using the elution buffer provided by the Ligation sequencing kit.

Regarding multiplexing experiments, we follow ONT recommendation and used two protocols: Native barcoding expansion 1-12 and 13-24 kit (ONT, catalog n°EXP-NBD104) and Native Barcoding lot 24 V14 (ONT, catalog n°SQK-NBD114.24). The first steps are similar than a singleplex experiment, from gDNA dephosphorylation step to Cas9 cleavage and dA-tailing step. Native barcodes were then ligated to the dA-tailed DNA, using blunt/TA ligase Master Mix (NEB), then followed by the adapter ligation step. As mentioned above, depending on which sequencing kit was used (V109 or V114), different adapter mix were used, either “AMII” buffer for V109 kit or “NA” buffer for V114 kit.

CRISPR/Cas9-targeted sequencing protocol was discontinued by ONT in 2024. Therefore, we adapted their protocol using similar existing products. For the first steps (gDNA dephosphorylation to dA- tailing), each ONT’s buffers and enzymes can be replaced by NEB’s products (Table 4). The adapter ligation step was performed following the Ligation Sequencing kit V114 protocol.

**Table 4.**
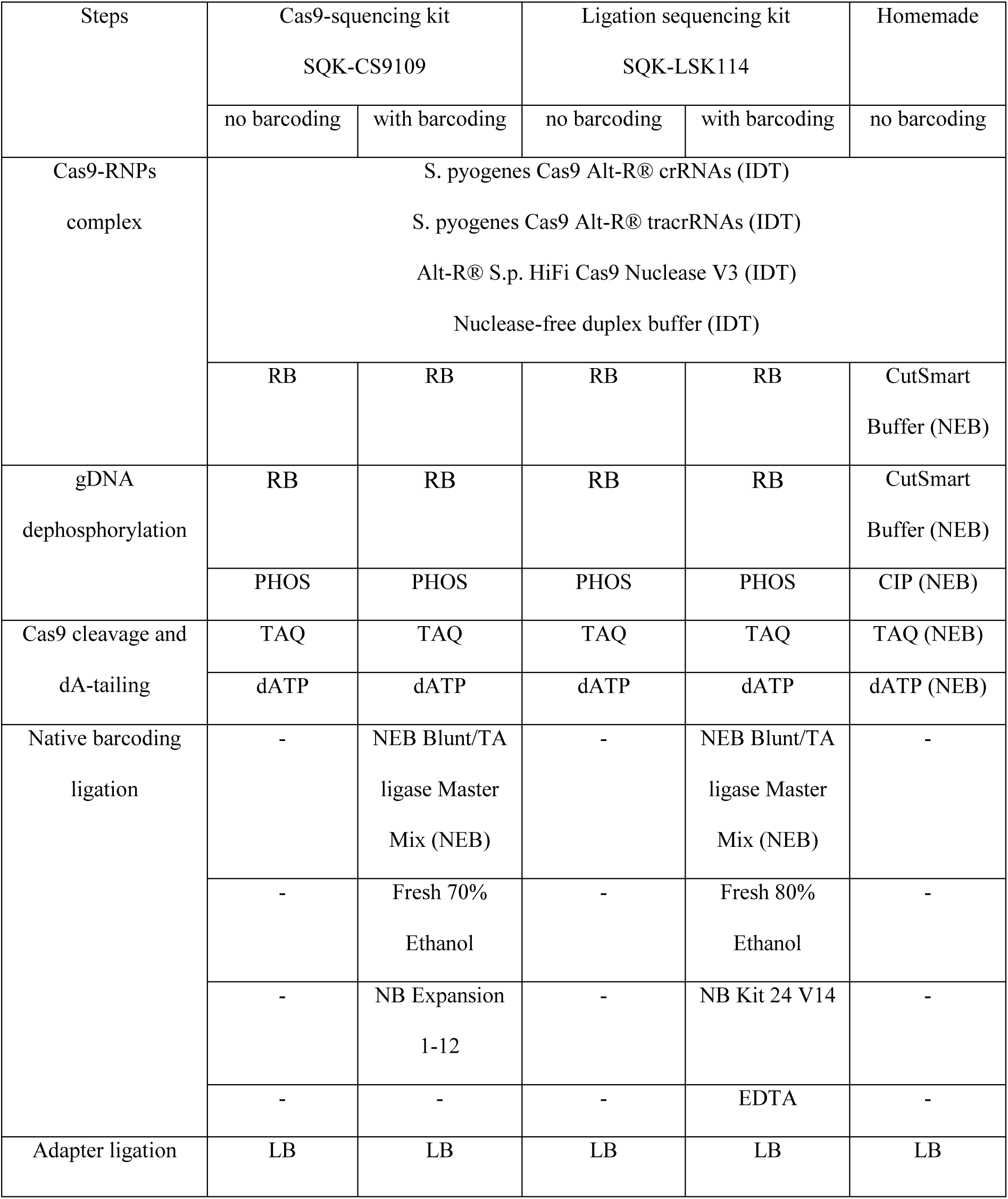
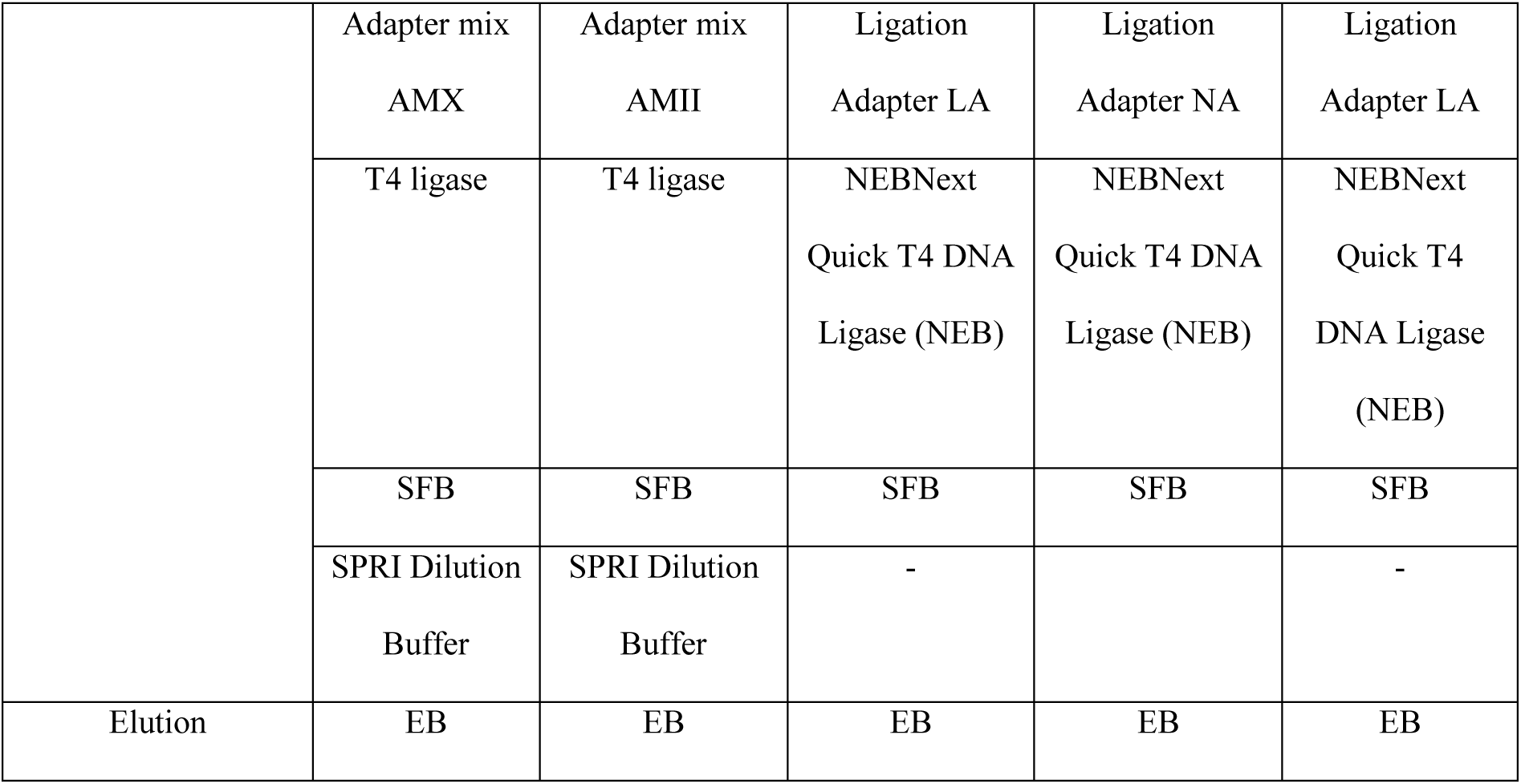
ONT long-read sequencing protocols. All reagents available using ONT ligation sequencing kit are referred with (ONT); other reagents are obtained through New England Biolabs (NEB) or Integrated DNA Technologies (IDT). RB – Reaction Buffer; PHOS – phosphatase; TAQ – Taq polymerase; NB – Native Barcoding; LB – ligation buffer; SFB – Short fragment buffer; EB – Elution Buffer

| Steps | Cas9-squencing kit<br>SQK-CS9109 |  | Ligation sequencing kit<br>SQK-LSK114 |  | Homemade |
| --- | --- | --- | --- | --- | --- |
|  | no barcoding | with barcoding | no barcoding | with barcoding | no barcoding |
| Cas9-RNPs<br>complex | S. pyogenes Cas9 Alt-R® crRNAs (IDT)<br>S. pyogenes Cas9 Alt-R® tracrRNAs (IDT)<br>Alt-R® S.p. HiFi Cas9 Nuclease V3 (IDT)<br>Nuclease-free duplex buffer (IDT) |  |  |  |  |
|  | RB | RB | RB | RB | CutSmart<br>Buffer (NEB) |
| gDNA<br>dephosphorylation | RB | RB | RB | RB | CutSmart<br>Buffer (NEB) |
|  | PHOS | PHOS | PHOS | PHOS | CIP (NEB) |
| Cas9 cleavage and<br>dA-tailing | TAQ | TAQ | TAQ | TAQ | TAQ (NEB) |
|  | dATP | dATP | dATP | dATP | dATP (NEB) |
| Native barcoding<br>ligation | - | NEB Blunt/TA<br>ligase Master<br>Mix (NEB) | - | NEB Blunt/TA<br>ligase Master<br>Mix (NEB) | - |
|  | - | Fresh 70%<br>Ethanol | - | Fresh 80%<br>Ethanol | - |
|  | - | NB Expansion<br>1-12 | - | NB Kit 24 V14 | - |
|  | - | - | - | EDTA | - |
| Adapter ligation | LB | LB | LB | LB | LB |
|  | Adapter mix<br>AMX | Adapter mix<br>AMII | Ligation<br>Adapter LA | Ligation<br>Adapter NA | Ligation<br>Adapter LA |
|  | T4 ligase | T4 ligase | NEBNext<br>Quick T4 DNA<br>Ligase (NEB) | NEBNext<br>Quick T4 DNA<br>Ligase (NEB) | NEBNext<br>Quick T4<br>DNA Ligase<br>(NEB) |
|  | SFB | SFB | SFB | SFB | SFB |
|  | SPRI Dilution<br>Buffer | SPRI Dilution<br>Buffer | - |  | - |
| Elution | EB | EB | EB | EB | EB |

Library quantification and quality controls were performed using the Qubit fluorometer (Fisher Scientific) and the TapeStation System with the Genomic DNA ScreenTape analysis kit (Agilent, catalog n°5067-5365). Libraries were sequenced using FLO-MIN106D (R9.4) or FLO-PRO114 (R10.4) flow cell using PromethION sequencer.

#### Adaptive sampling

A quantity of 3.5 µg of double-stranded DNA assayed by Qubit fluorometer was fragmented with g-Covaris TUBE (Covaris, catalog n°520079) to a size of approximately 12 kb using an Eppendorf 5418 centrifuge (Merck, Saint-Quentin-Fallavier, France) at a speed of 14,000 rpm for 1min following the covaris protocol. This size was assessed by TapeStation using Genomic ScreenTape (Agilent). The library was prepared using SQK-LSK114 ligation kit from ONT and associated NEB enzymes according to the manufacturer’s recommendations (genomic-dna-by-ligation-sqk-lsk114- GDE_9161_v114_revP_29Jun2022-promethion.pdf). A quantity of 180 fmoles is obtained at the end of the preparation and loaded onto the PromethION FLO-PRO114M flow cell (ONT). Sequencing with enrichment by adaptive sampling is performed for 48h total on PromethION P24 (PRO-SEQ024 and PRO-PRCA100). The enrichment bed file targets the short arm of chromosome 19 (Chr19: 26,200,001- 58,617,616) with the hg38 reference genome in fasta format. Raw data pod5 were generated using ONT MinKnow software (v24.06.15) and basecalling of fastq files was performed in super precision mode (Sup AC) directly on the PromethION P24 using Dorado software (version 7.4.14). This sequencing generated 23Gb of raw data.

#### Whole genome long-read sequencing

Double stranded DNA was sheared using Covaris g-TUBE (Covaris) at an approximated size of 20kb, by centrifuging g-TUBE at 4200 rpm for 1 minute, according to the manufacturer’s recommendations. DNA fragment size was evaluated using Genomic DNA ScreenTape Analysis (Agilent). Library preparation was then performed according to the ligation sequencing kit V114 (SQK-LSK114, ONT). Half of the library (20 fmoles) were loaded into the PromethION flow cell FLO-PRO114M. After 30h, the flow cell was washed using a wash kit (EXP- WSH004, ONT) and the rest of the library preparation was then loaded into the flow cell and sequenced for 96 hours total on PromethION sequencer. Raw data (Fast5 files) were generated using Minknow software (v23.04.06) and basecalling if fastq files was performed in SupAC mode directly onto the PromothION sequencer server using Dorado BC (v6.5.7).

### Bioinformatics analyses

FAST5 files were processed using MinKnow software, as well as two open-source basecallers, Dorado and Bonito BC employing the high accuracy (HAC) model. Each version and neural network used during this study is specified in Table 5. MinKnow is proprietary software, and the first samples could not be analyzed with the same version or model used for the #16814 sample. We took this important parameter into consideration and used the standalone version of Dorado BC (v5.3) to compare patients and discuss the differences of performances between basecallers. FASTQ files were aligned to the T2T genome utilizing Minimap2. Minimap2 was run with the recommended parameter for ONT long reads (-a –x map_ont). Reads overlapping genes of interest were extracted via Samtools. To identify reads spanning the repeat region, local alignment was performed with adjacent regions, each spanning 100 bases before and after the repeat site. All retained reads were oriented in the same direction. The repeat region was precisely located by identifying the two 100-base regions flanking the repeat, as taken from the T2T consensus sequences. Reads exhibiting over 90% identity with the consensus on each flanking region. The count of repeats was determined by dividing the total number of bases in between the 2 flanking regions by the length of the repeat motif. We also used Straglr on the same set of reads. Straglr was modified to focus on clinical tandem repeat (TR) (https://github.com/philres/straglr) with the recommended parameters from the website. Methylation status was assessed using Modkit on the same reads after transformation to POD5. Methylation was called using Dorado (v5.3) with the modified-bases model (Dorado basecaller HAC for 5mCG, 5hmCG and 6mA) with DNA models available at https://github.com/nanoporetech/dorado. Reads were stratified into ’expanded’ and ’non-expanded’ subgroups based on the largest gap in length observed in the sorted list of read lengths. Visualization used R and Gviz. Files are available on GitHub (https://github.com/PYB-SU/STR).

**Table 5.**
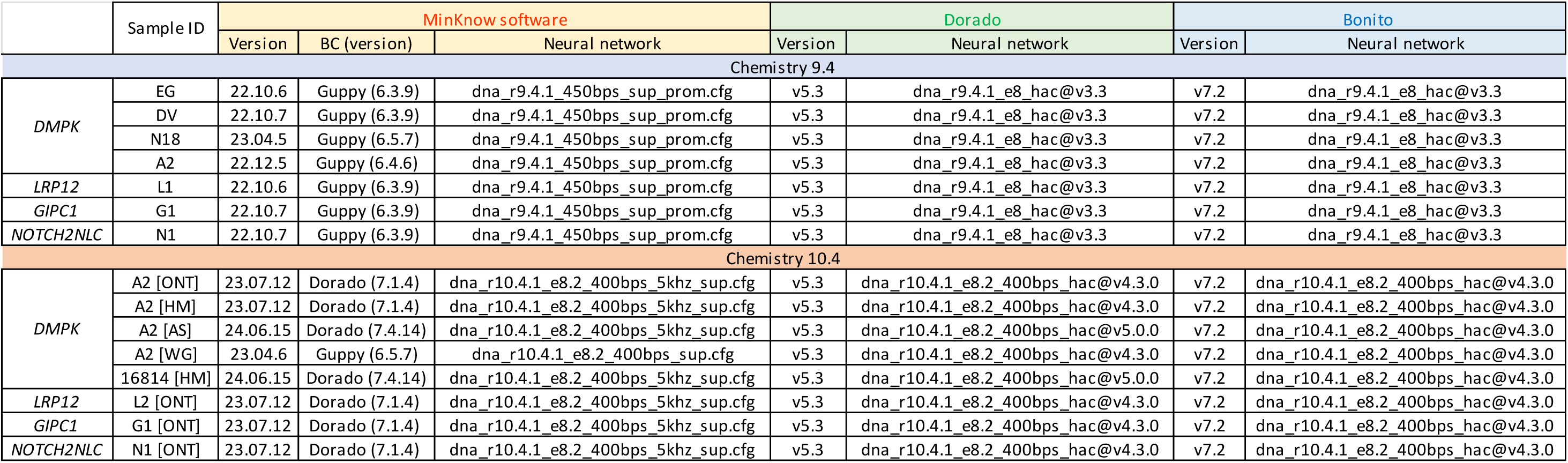
Basecaller versions and associated neural networks. Three basecalling strategies were tested: MinKnow (yellow), Dorado (green) and Bonito (blue). For each sample, basecaller and associated neural network versions are referenced.

## ETHICAL STATEMENT

### Patients’ consent

All patient participating in this study signed a written informed consent allowing us to use their biological material (here DNA) in a research setting.

## CODE AVAILABILITY

Bioinformatical pipelines used in this study are open-sourced and available on GitHub. Hyperlinks are available in the method section “Bioinformatics analyses”.

### CRediT AUTHOR STATEMENT

Conceptualization – **Louise Benarroch** and **Stéphanie Tomé**

Formal analysis – **Badreddine Mohand-Oumoussa**, **Hélène Madry**, **Pierre-Yves Boëlle**, **Karim Labrèche**

Funding acquisition – **Gisèle Bonne** and **Stéphanie Tomé**

Investigations – **Louise Benarroch**, **Stéphanie Tomé**, **Pierre-Yves Boëlle**

Methodology – **Louise Benarroch**, **Stéphanie Tomé**, **Badreddine Mohand-Oumoussa**, **Hélène Madry** and **Valeriia Gorbunova**.

Ressources – **Nobuyuki Eura**, **Ichizo Nishino**, **Tanya Stojkovic**, **Guillaume Bassez**, **Gisèle Bonne**

Writing, original draft – **Louise Benarroch** and **Stéphanie Tomé**

Writing, review and editing – **Louise Benarroch**, **Stéphanie Tomé**, **Pierre-Yves Boëlle**, **Gisèle Bonne**, **Geneviève Gourdon**

## ACKNOWLEDGEMENTS

This work was funded by Association Institut de Myologie (AIM, Paris France), INSERM (Paris, France) and Sorbonne Université (Paris, France). Louise Benarroch was supported by Solve-RD project which has received funding from the European Union’s Horizon 2020 research and innovation program under grant agreement No 779257 (PI: Gisèle Bonne) and Muscular Dystrophy United Kingdom (MDUK) grant No. 18GROI-PG24-0140-2 (PI: Gisèle Bonne, United Kingdom). This work was also supported by a grant from Fondation Maladies Rares N°GenOmics (2023)_041704 (PI: Stéphanie Tomé, France).

## DECLARATION OF AI AND AI-ASSOCIATED TECHNOLOGY

During the preparation of this work the authors used ‘Deepl Translator’ and ‘Chat GPT’ in order to translate from French to English and to correct errors in synthax, grammar and conjugation. After using these tools, the authors reviewed and edited the content as needed and take full responsibility for the content of the publication.

