## Supplementary figures for "Comparative analysis of CRISPR/Cas9-targeted nanopore sequencing approaches in repeat expansion disorders"

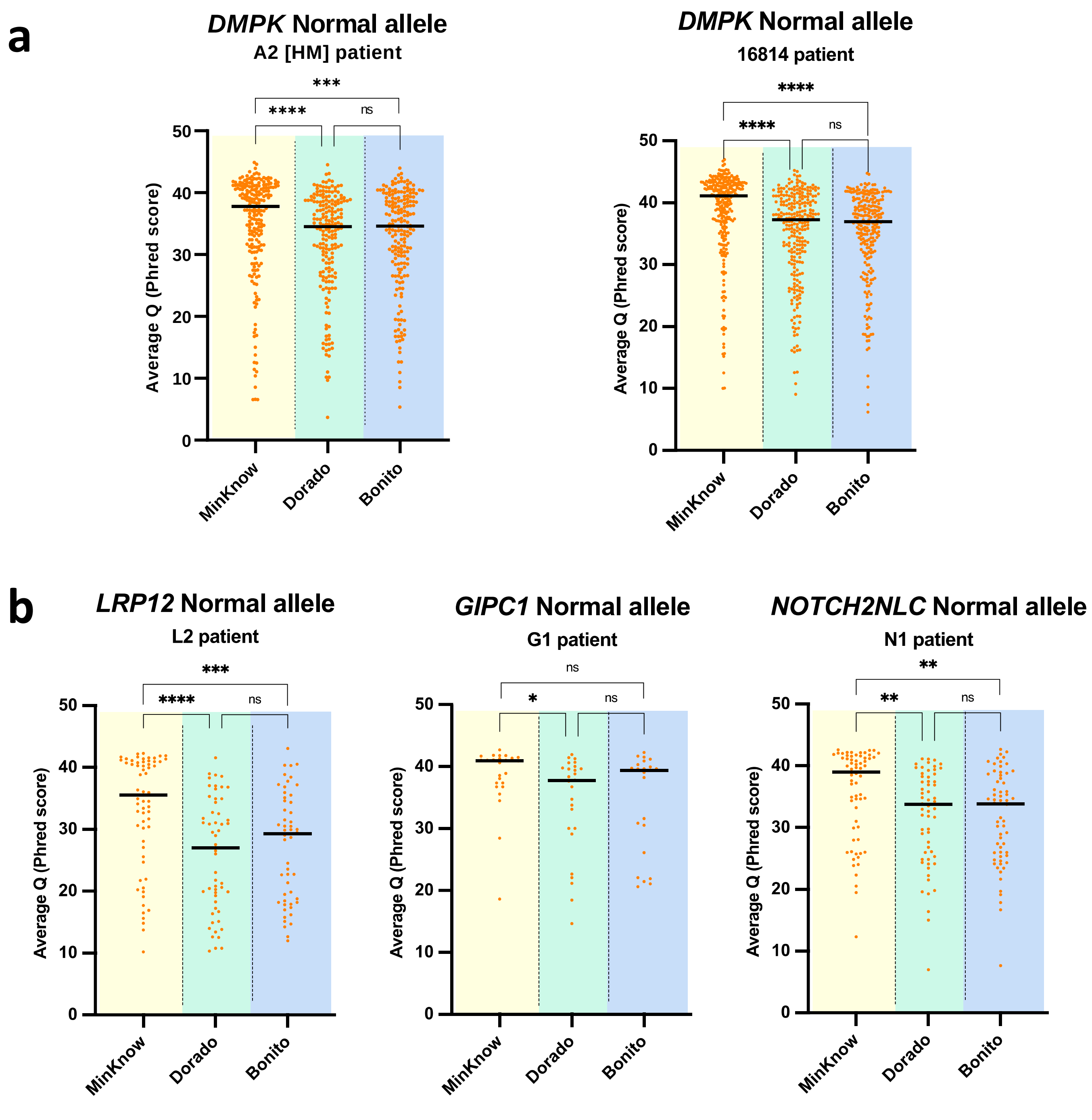


**Figure S1.** Normal allele average quality. Basecallers' performance comparison using R10.4 chemistry at **(a)** *DMPK* locus with A2 [HM] (left panel) and #16814 (right panel) patients, and **(b)** OPDM loci: *LRP12* (left panel), *GIPC1* (middle panel) and *NOTCH2NLC* (right panel). Kruskal-Wallis test, ns – not significant; [*] p-value < 0.05; [**] p-value < 0.01; [***] p-value < 0.001 and [****] p-value < 0.0001

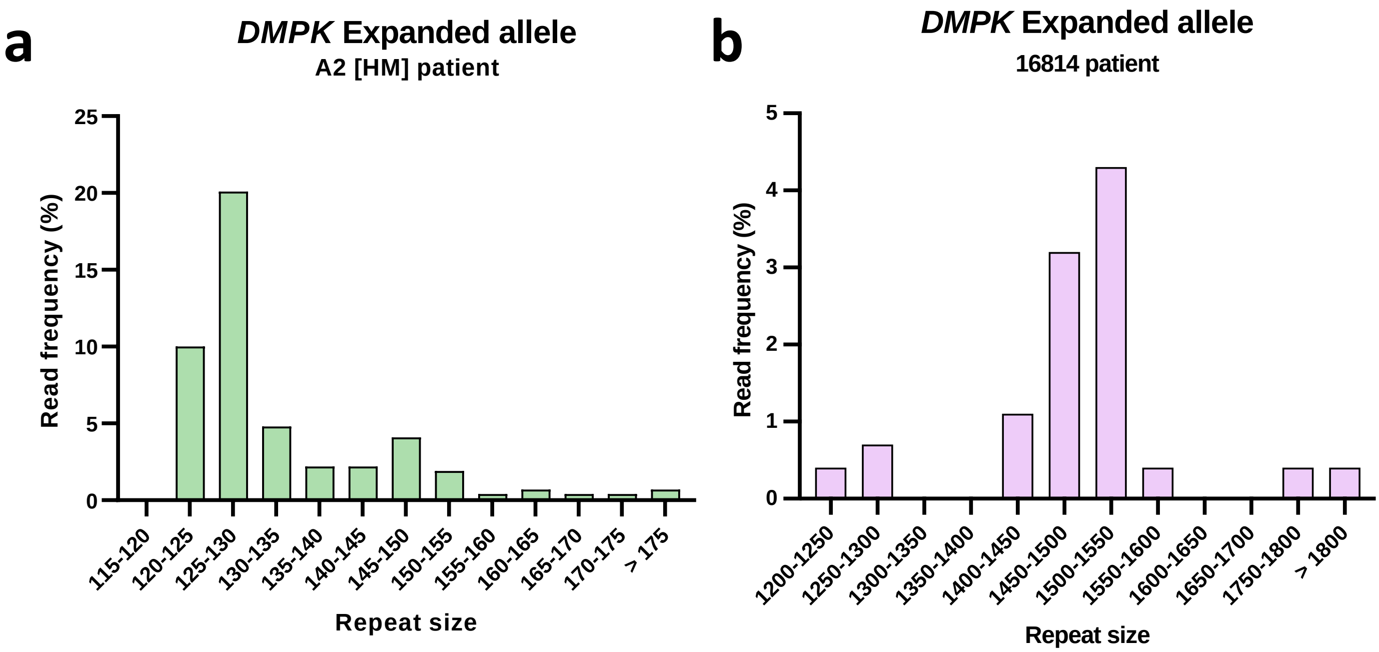


**Figure S2.** Read length distribution of *DMPK* expanded allele. **(a)** A2 [HM] patient and **(b)** #16814 patient. The x-axis displays the length of the repeats detected (number of repeats, 3bp per repeat). The y-axis displays read frequency calculated as a ratio of number of Q20 reads on total number of reads.

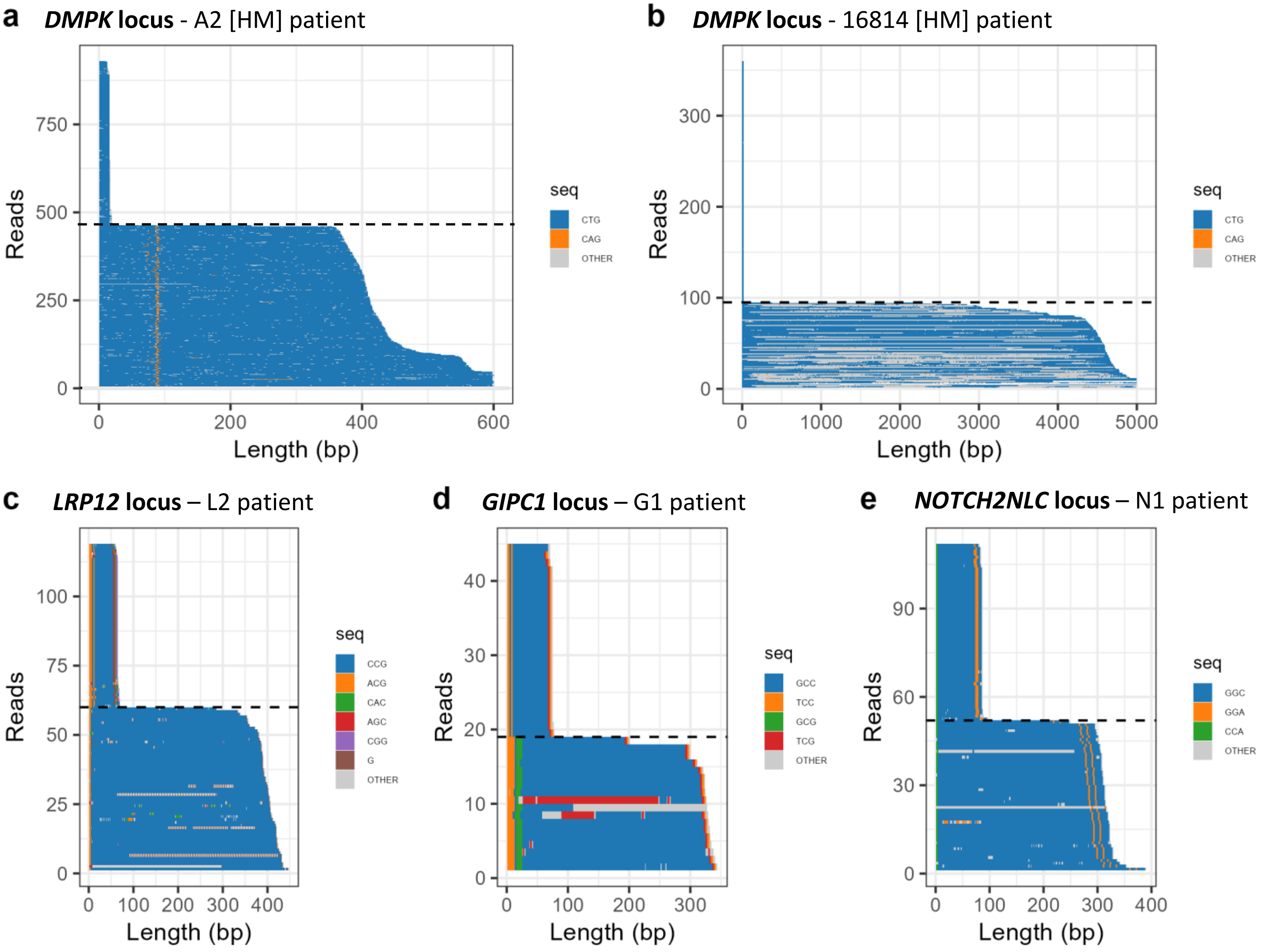


**Figure S3.** Waterfall plots highlighting the expansion structures at **(a-b)** DM1 locus and **(c-e)** OPDM loci, with a phred score > 10. For each waterfall, the x-axis shows the repeat expansion length in base pairs. The y-axis displays all the reads obtained after sequencing. The reference codon **(a-b)** CTG, **(c)** CCG, **(d)** GCC or **(e)** GGC is represented in blue; different colours are used to highlight codons within the repeat sequence that are not the reference codon, within the expansion. Artefactual interruptions are highlighted in grey.

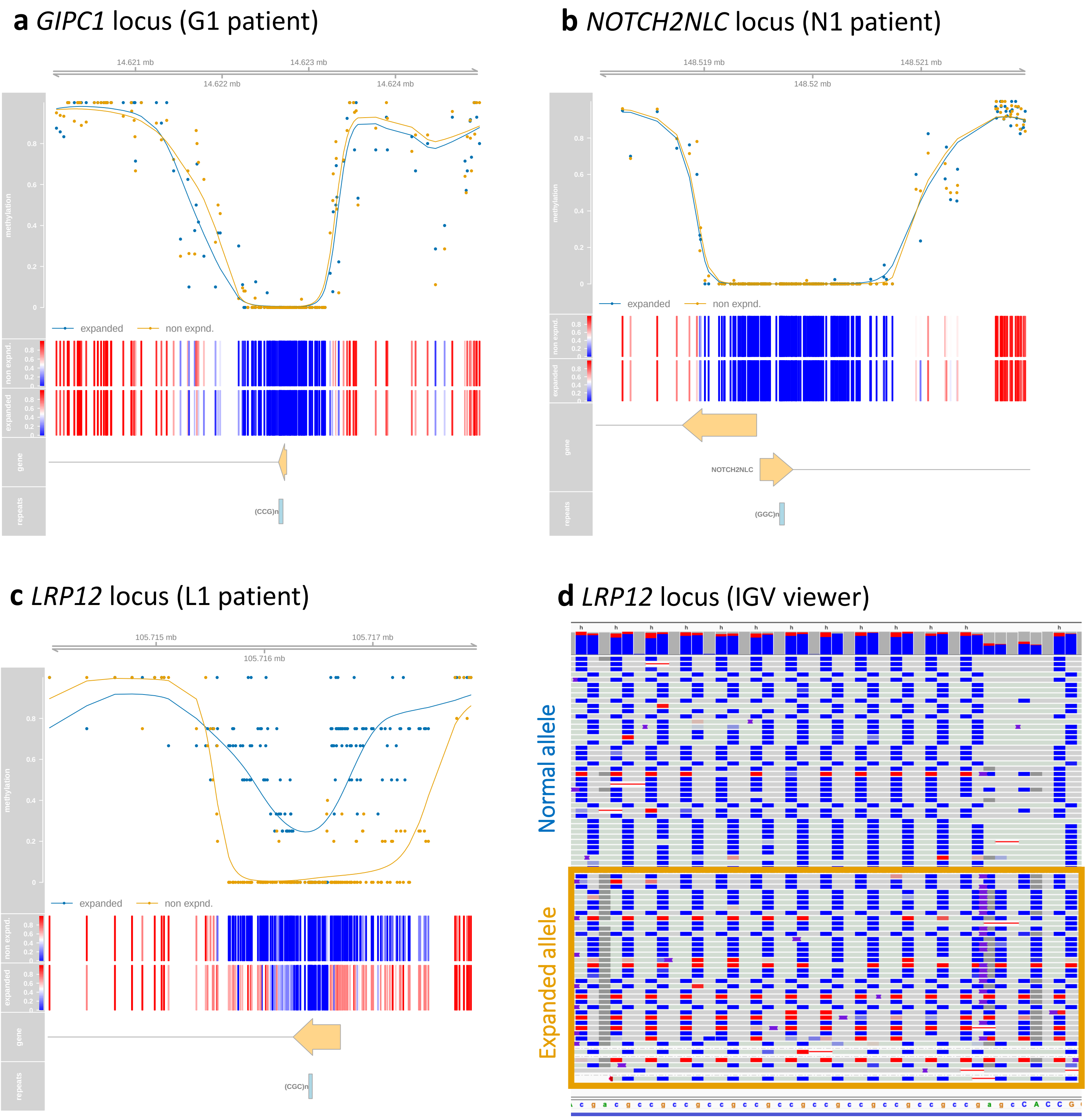


**Figure S4.** 5-methylcytosine (5mC) methylation levels at **(a)** *GIPC1* (G1 patient) and **(b)** *NOTCH2NLC* (N1 patient) and **(c-d)** *LRP12* (L1 patient, R9.4 chemistry) loci. Each graph displays (a-c), from top to bottom, the genomic localization; the methylation levels where each dot represent one read on the normal allele (yellow) or the expanded allele (blue); heatmap of methylation levels; gene(s) located in the region of interest; and localization of the repeats. **(d)** IGV screenshot of the *LRP12* expanded region for L1 patient, displaying the methylation status of each read. The normal allele on top and the expanded allele at the bottom (surrounded by a yellow frame). Each line represents one read. Unmethylated sites are colored in blue, methylated sites are colored in red.

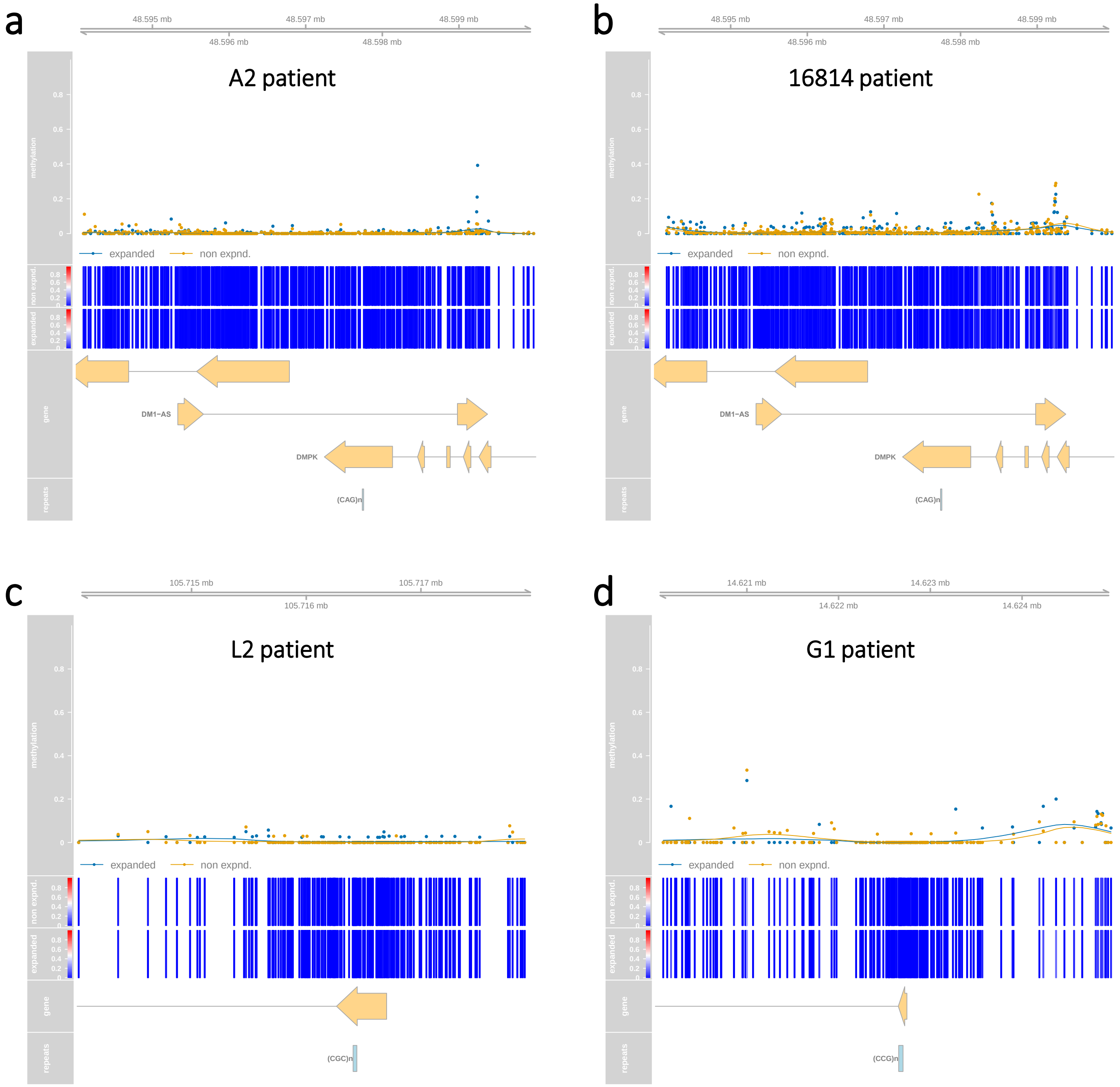


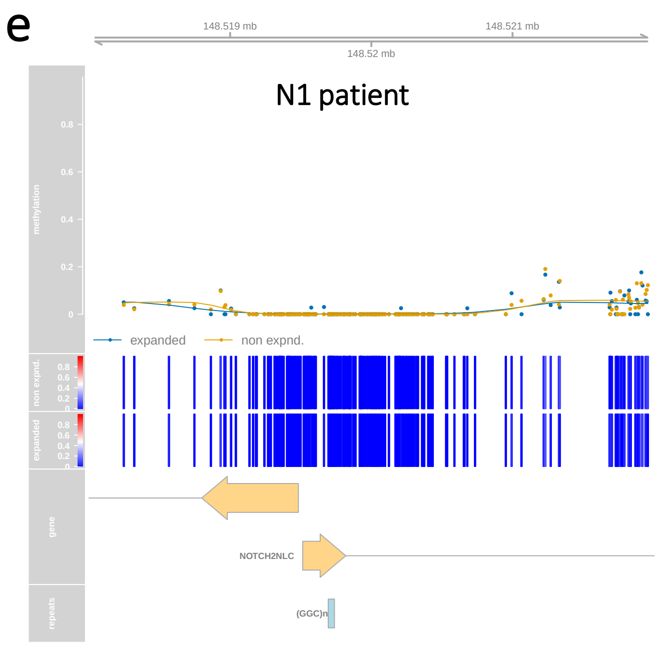


**Figure S5.** 5-hydroxymethylcytosine (5hmC) methylation levels of **(a-b)** DM1 and **(c-e)** OPDM loci. Methylation profile for **(a)** A2 [HM], **(b)** #16814, **(c)** L2, **(d)** G1 patient and **(e)** N1 patients. Each graph displays, from top to bottom, the genomic localization; the methylation levels where each dot represent one read on the normal allele (yellow) or the expanded allele (blue); heatmap of methylation levels; gene(s) located in the region of interest; and localization of the repeats.

| Norm. allele (10.4) | Sample ID | MinKnow | | | Dorado | | | Bonito | | |
|---|---|---|---|---|---|---|---|---|---|---|
| | | AvgQ (all) | %Q20 | AvgQ (Q20) | AvgQ (all) | %Q20 | AvgQ (Q20) | AvgQ (all) | %Q20 | AvgQ (Q20) |
| *DMPK* | A2 [ONT] | 35.0 | 93% | 36.8 | 32.2 | 89% | 34.4 | 32.6 | 88% | 35.0 |
| | A2 [HM] | 36.2 | 93% | 37.9 | 33.2 | 93% | 34.8 | 33.5 | 91% | 35.6 |
| | A2 [AS] | 24.6 | 67% | 29.2 | 32.4 | 88% | 35.3 | 30.3 | 81% | 34.2 |
| | A2 [WG] | 30.9 | 100% | 30.9 | 31.3 | 92% | 32.5 | 33.2 | 83% | 39.9 |
| | 16814 [HM] | 38.5 | 95% | 39.5 | 34.9 | 94% | 36.1 | 35.0 | 94% | 36.3 |
| *LRP12* | L2 [ONT] | 33.0 | 86% | 35.9 | 25.8 | 67% | 31.1 | 27.2 | 67% | 32.3 |
| *GIPC1* | G1 [ONT] | 38.4 | 96% | 39.2 | 34.0 | 92% | 35.4 | 34.3 | 100% | 34.3 |
| *NOTCH2NLC* | N1 [ONT] | 35.6 | 97% | 36.2 | 31.7 | 90% | 33.4 | 31.6 | 92% | 32.9 |

**Table S1.** Basecaller comparison on average quality of all reads of the normal allele. Three basecallers were tested: MinKnow (yellow), Dorado (green) and Bonito (blue). AvgQ - Average quality (phred score); %Q20 - percentage of Q20 reads on total expanded allele reads.
