## Supplementary file for "Comparative analysis of CRISPR/Cas9-targeted nanopore sequencing approaches in repeat expansion disorders"

**File S1.** OPDM loci sequences.
For all OPDM genes, the normal and expanded sequence have been extracted using Guppy basecaller. The underlined codon was used to align all reads. Color code used in the sequence match color coding in waterfall plots.

*LRP12* loci / L2 patient (R10.4)

**Normal allele**

5'- AGG CCA TAA CCA CAG CAG ATG GAG AGA GAG AGA GGA GGA GAC GGA GGA GGA GGG
AGG AGG AGA AGC TGG AGG **<u>TAG</u>** **ACG ACG CCG ACG CCG CCG CCG CCG CCG CCG CCG CCG**
**CCG CCG CCG CCG CCG CCG AGC CAC CGG** CTG CTC CCT GCG CTC TCC GCG GCT GCG GGA
GGG GGA AGG GAG GGG CCG CCG CCG CCC GCG CGC GCT CCC TCC -3'

**Expanded allele**

5'- GCG ACA GGC CAT AAC CAC AGC AGA TGG AGA GAG AGA GGA GGA GAC GGA GGA GGA GGG
AGG AGG AGC TGG AGG **<u>TAG</u>** **ACG CCG CCG CCG CCG CCG CCG CCG CCG CCG CCG CCG CCG CCG**
**CCG CCG CCG CCG CCG CCG CCG CCG CCG CCG CCG CCG CCG CCG CCG CCG CCG CCG CCG CCG**
**CCG CCG CCG CCG CCG CCG CCG CCG CCG CCG CCG CCG CCG CCG CCG CCG CCG CCG CCG CCG**
**CCG CCG CCG CCG CCG CCG CCG CCG CCG CCG CCG CCG CCG CCG CCG CCG CCG CCG CCG CCG**
**CCG CCG CCG CCG CCG CCG CCG CCG CCG CCG CCG CCG CCG CCG CCG CCG CCG CCG CCG CCG**
**CCG CCG CCG CCG CCG CCG CCG CCG CCG CCG CCG CCG CCG CCG CCG CCG CCG CCG CCG CCG**
**CCG CCG CCG CCG CCG CCG CCG CCG CCG CCG CCG CCG CCG CCG CCG CCG CCG CCG CCG CCG**
**CCG G** CTG CTC CCT GCG CTC TCC GCG GCT GCG GGA GGG GGA AGG GAG GGG CCG CCG
CCG CCC GCG CGC GCT CCC TCC -3'

*LRP12* loci / L1 patient (R9.4)

**Normal allele**

5'- CCA TAA CCA CAG CAG ATG GAG AGA GAG GGA GAC GGA GGA GGA GGG AGG AGA
AGC TGG AGG **<u>TAG</u>** **ACG ACG CCG CCG CCG CCG CCG CCG CCG CCG CCG CCG CCG AGC CAC**
**CGG** CTG CTC CCT GCG CTC TCC GCG GCT GCG GGA GGG GGA AGG GAG GGG CCG CCG CCG
CCC GCG CGC GCT CCC TCC -3'

**Expanded allele**

5'- GCG ACA GGC CAT AAC CAC AGC AGA TGG AGA GAG AGA GGA GGA GAC GGA GGA GGA
GGG AGG AGG AGC TGG AGG **<u>TAG</u>** **ACG CCG CCG CCG CCG CCG CCG CCG CCG CCG CCG CCG**
**CCG CCG CCG CCG CCG CCG CCG CCG CCG CCG CCG CCG CCG CCG CCG CCG CCG CCG CCG CCG**
**CCG CCG CCG CCG CCG CCG CCG CCG CCG CCG CCG CCG CCG CCG CCG CCG CCG CCG CCG CCG**
**CCG CCG CCG CCG CCG CCG CCG CCG CCG CCG CCG CCG CCG CCG CCG CCG CCG CCG CCG CCG**
**CCG CCG CCG CCG CCG CCG CCG CCG CCG CCG CCG CCG CCG CCG CCG CCG CCG CCG CCG CCG**
**CCG CCG CCG CCG CCG CCG CCG CCG CCG CCG CCG CCG CCG CCG CCG CCG CCG CCG CCG CCG**
**CCG CCG CCG CCG CCG CCG CCG CCG CCG CCG CCG CCG CCG CCG CCG CCG CCG CCG CCG CCG**
**CCG CCG CCG CCG CCG CCG CCG CCG CCG CCG CCG CCG CCG G** CTG CTC CCT GCG CTC TCC
GCG GCT GCG GGA GAA GGG AGG GGA GCC GCC GCC GCC CGC GCG CGC TCC CTC CTC CGT
CCT CCC TCC GGC TCG CCT -3'

*GIPC1* loci / G1 patient

**Normal allele**

5'- CCG CGG CGA CGC CCT CTC CGG GCC AGA CCT CAA ACG CCT CCC GGC TCG GCC CAC
GCG TGC **CCC** TCC GCC TCC GCC GCC GCC GCC GCC GCC GCC GCC GCC GCC GCC GCC GCC
GCC GCC GCC GCC GCC GCC TCG TCC ACT CAC CGG AGA CGC TCT TCC TAC ACA GAC GAG
GCC CCC GGA GCC CCC AGC GCC CGG ACG -3'

**Expanded allele**

5'- CCG CGG CGA CGC CCT CTC CGG GCC AGA CCT CAA ACG CCT CCC GGC TCG GCC CAC
GCG TGC **CCC** TCC TCC TCC TCC GCC GCG GCG GCG GCC GCC GCC GCC GCC GCC GCC GCC
GCC GCC GCC GCC GCC GCC GCC GCC GCC GCC GCC GCC GCC GCC GCC GCC GCC GCC GCC
GCC GCC GCC GCC GCC GCC GCC GCC GCC GCC GCC GCC GCC GCC GCC GCC GCC GCC GCC
GCC GCC GCC GCC GCC GCC GCC GCC GCC GCC GCC GCC GCC GCC GCC GCC GCC GCC GCC
GCC GCC GCC GCC GCC GCC GCC GCC GCC GCC GCC GCC GCC GCC GCC GCC GCC GCC GCC
GCC GCC GCC GCC GCC GCC GCC GCC GCC GCC GCC GCC GCC GCC TCG TCC ACT CAC CGG
AGA CGC TCT TCC TAC ACA GAC GAG GCC CCC GGA GCC CCC AGC GCC CGG ACG

*NOTCH2NLC* loci / N1 patient

**Normal allele**

5'- GAG AGT GGG CTC CTC TAT CGG GAC CCC CTC CCC ATG TGG ATC **TGC** CCA GGC GGC
GGC GGC GGC GGC GGC GGC GGC GGC GGC GGC GGC GGC GGC GGC GGC GGC GGC GGC GGC
GGC GGA GGA GGC GGC GAC CGA GAA GAT GCC CGC CCT GCG CCG CTC TGC TGT -3'

**Expanded allele**

5'- GAG AGT GGG CTC CTC TAT CGG GAC CCC CTC CCC ATG TGG ATC **TGC** CCA GGC GGC
GGC GGC GGC GGC GGC GGC GGC GGC GGC GGC GGC GGC GGC GGC GGC GGC GGC GGC GGC
GGC GGC GGC GGC GGC GGC GGC GGC GGC GGC GGC GGC GGC GGC GGC GGC GGC GGC GGC
GGC GGC GGC GGC GGC GGC GGC GGC GGC GGC GGC GGC GGC GGC GGC GGC GGC GGC GGC
GGC GGC GGC GGC GGC GGC GGC GGC GGC GGC GGC GGC GGC GGC GGC GGC GGC GGC GGC
GGC GGC GGC GGC GGC GGC GGC GGC GGC GGC GGC GGC GGC GGC GGC GGA GGC GGC GGC
GGA GGC GGC GGC GGC GGC GAC CGA GAA GAT GCC CGC CCT GCG CCG CTC TGC TGT -3'
